# Genomic and structural insights into Jyvaskylavirus, the first giant virus isolated from Finland

**DOI:** 10.1101/2024.10.16.618641

**Authors:** Gabriel Magno de Freitas Almeida, Iker Arriaga, Bruna Luiza de Azevedo, Miika Leppänen, Jonatas Santos Abrahao, Julien Andreani, Davide Zabeo, Janne Ravantti, Nicola GA Abrescia, Lotta-Riina Sundberg

**Affiliations:** The Norwegian College of Fishery Science, Faculty of Biosciences, Fisheries and Economics, UiT □ The Arctic University of Norway, Tromsø, Norway; Structure and Cell Biology of Viruses Lab, CIC bioGUNE, Basque Research and Technology Alliance (BRTA), Derio, Spain; Universidade Federal de Minas Gerais, Institute of Biological Sciences, Department of Microbiology, Belo Horizonte, Minas Gerais, Brazil; University of Jyväskylä, Department of Biological and Environmental Science and Nanoscience Center, Jyväskylä, Finland; Aix Marseille Univ, MEPHI, Marseille, France; IHU-Méditerranée infection, Marseille, France; Diamond Light Source, Harwell Science and Innovation Campus, Didcot, UK; University of Helsinki, Molecular and Integrative Biosciences Research Programme, Helsinki, Finland; Ikerbasque, Basque Foundation for Science, Bilbao, Spain

## Abstract

Giant viruses of protists are a diverse and likely ubiquitous group of organisms. Here, we describe Jyvaskylavirus, the first giant virus isolated from Finland. This clade B marseillevirus was found in *Acanthamoeba castellanii* from a composting soil sample in Jyväskylä, Central Finland. Its genome shares similarities with other marseilleviruses. Helium ion microscopy and electron microscopy of infected cells unraveled stages of the Jyvaskylavirus lifecycle. We reconstructed the Jyvaskylavirus particle to 6.3 Å resolution using cryo-EM. The ∼2,500 Å diameter virion displays structural similarities to other *Marseilleviridae* giant viruses. The capsid comprises of 9,240 copies of the major capsid protein ORF184, which possesses a double jellyroll fold arranged in trimers forming pseudo-hexameric capsomers. Below the capsid shell, the internal membrane vesicle encloses the genome. Through cross-structural and -sequence comparisons with other *Marseilleviridae* using AI-based software in model building and prediction, we elucidated ORF142 as the penton protein, which plugs the twelve vertices of the capsid. Five additional ORFs were identified, with models predicted and fitted into densities that either cap the capsomers externally or stabilize them internally. The isolation of Jyvaskylavirus suggests that these viruses may be widespread in the boreal environment and provide structural insights extendable to other marseilleviruses.

## INTRODUCTION

Viruses defy paradigms of classical biology and are agents of change and innovation for biological research. In 2003 researchers were surprised by the description of the *Acanthamoeba polyphaga mimivirus* (APMV) (**La Scola et al 2003**). Hidden in plain sight for over a century of microbiology research, APMV and other giant viruses were evading discovery for three main reasons: their large size traps them in filters commonly used in virological works, their structure and size makes them visible by Gram-staining, and their protist hosts are not as well studied as other host species (**Colson et al 2017; Queiroz et al 2022**). Common features of their double-stranded DNA genomes, similarity in a few core genes, relative independence of host transcription machinery, partial or complete cytoplasmic replication and formation of viral factories include giant viruses in the nucleo-cytoplasmic large DNA viruses (NCLDVs) group (**Iyer et al 2001; Boyer et al 2010**). Recently the International Committee of Taxonomy of Viruses (ICTV) classified giant viruses as members of the phylum *Nucleocytoviricota* of the kingdom *Bamfordvirae* in the realm *Varidnaviria* (**Walker et al, 2019**; **Simmonds et al 2023**). Isolation efforts and metagenomic data in the last two decades have revealed that giant viruses are ubiquitous in the environment (**Moniruzzaman et al 2020; Schulz et al 2020**). Giant virus particles or their DNA have been found from the Antarctica to the Siberian permafrost, in deep-sea sediments and many other sources including urban environments and diverse sample types (**Andrade et al 2018**; **Legendre et al 2015**; **Backstrom et al 2021**; **Colson et al 2017**). To support the sharp rise in metagenomic studies, microbial isolation and characterization are welcomed, and needed for deeper understanding of these entities and the viral world. Isolation of giant viruses revealed the intriguing mosaicism of Marseillevirus, the complex structure and mysterious large genome of Pandoravirus, the remarkable structure and translational potential of Tupanvirus, and many other characteristics of these organisms that have changed our view on the concept of viruses and their evolution (**Boyer et al 2009**; **Phillippe et al 2013**; **Abrahao et al 2018**). It is expected that many new insights will emerge as more isolates are found (**Aherfi et al 2016; Colson et al 2017**). Increased effort in isolating giant viruses might also bring new discoveries and help in understanding their distribution and importance worldwide. Isolation of viruses allows for in-depth structural studies using their whole virions.

An increased number of icosahedral NCLVD, thanks to the advances in cryo-EM, are now being targeted for structural analysis despite the challenges that their very large dimensions pose (> 1,500 Å diameter). Cryo-EM structures for African swine fever virus, *Aureococcus anophagefferens* virus, *Cafeteria roenbergensis* virus, Faustovirus, Marseillevirus, Medusavirus, Pacmanvirus, Paramecium bursaria Chlorella virus 1, *Phaeocystis pouchetii* virus, and more recently for Melbournevirus have elucidated their complex architecture, major capsid protein fold, and assembly organization (**Andreani et al 2017; Andres et al 2020; Burton-Smith et al 2021; Gann et al 2020; Klose et al 2016; Okamoto et al 2018; Shao et al 2022; Wang et al 2019; Watanabe et al 2022; Xiao et al 2017; Yan et al 2005**). Another example of the importance of the combination of isolation and structural studies is the description of FLiP, a missing link between ssDNA and dsDNA viruses in Finland (**Laanto et al 2017)**. However, FLiP is a bacteriophage and there have been no previous studies on the isolation of giant viruses in the Finnish boreal ecosystem or have their structural characterization been described so far.

Microbial ecology and virus-host interactions are still poorly studied in many environments, including boreal ecosystems. Here, we report the occurrence of giant viruses in Central Finland and the characterization of Jyvaskylavirus, a new virus belonging to the clade B of the *Marseilleviridae* family as determined by genome analysis. This virus was isolated from a composting soil sample in the city of Jyväskylä and represents the northernmost marseillevirus known to date. Other marseilleviruses from the northern hemisphere were found in France, India, Japan, Algeria and Senegal only (**Sahmi-Bounsiar et al 2021**), while closer giant viruses are either an uncharacterized Swedish cedratvirus (**Kordel et al 2021**) or a few microalgae-infecting mimivirus-like and phycodnaviruses-like isolates from Norway (**Catsberg et al 2002; Sandaa et al 2001; Johannessen et al 2015**). We used helium ion microscopy and transmission electron microscopy to visualize the early attachment events of the virus to its amoebal host (*Acanthamoeba castellanii*) and the infected cell. Using cryo-EM, we also determined the three-dimensional (3D) structure of Jyvaskylavirus at a resolution of 6.3 Å. Cross-structural and sequence comparison allowed us to identify five proteins that compose the capsid essential for assembly, with their corresponding models reliably placed into density.

Jyvaskylavirus description is the first step in unveiling the diversity of giant viruses from Finland and from the Nordic Countries, exemplifying that these viruses are also present in the boreal ecosystems with a still unknown role for microbial ecology.

## RESULTS

### Giant viruses are present in Finland

During the summer 2019 a preliminary isolation attempt of local amoebas and giant viruses using samples collected in Central Finland hinted at the presence of these viruses in Finnish samples (**Supplementary Material and Supplementary Figure 1**). Although viral-like particles were seen, working with the local protist host presented challenges that initially hindered the virus characterization. We followed up this preliminary screening with a larger isolation effort using three reference host strains: *Acanthamoeba castellanii*, *A. polyphaga* and *Vermamoeba vermiformis*. Ninety-six environmental samples collected in Central Finland were tested using the three different amoebal hosts. Of these samples, ten had viral presence confirmed by the appearance of cytophatic effect (CPE) in cultures of *A. castellanii* followed by visualization of negative stained viral-like particles by electron microscopy (10.41% isolation success in *A. castellanii*). However, some of the samples belonged to the same sample group and viral morphology was similar between some of the isolates. Adjusting for sample type and unique viral morphologies found, we have then tested 53 unique sample types and found three distinct viral morphologies (5.66% isolation success in *A. castellanii*) (see **Supplementary Table 1**).

Some of the CPE-positive samples contained mixed viral populations. Samples collected from experimental aquaria had a mixture of three different viral morphotypes with dimensions ranging from approximately 200 to 300 nm side-to-side, highlighting a previously unexplored diversity of giant viruses in this setting (**Figure 1A-C**). One sample from a recirculating aquaculture tank also possessed the diamond-shaped and loose-capsid morphologies seen in the aquaria samples. The high amount of nutrients in the fish aquaria or tanks might have enriched for protozoa and thus favoured our giant virus isolation, in a pattern similar to the original methodology for isolating giant viruses (**Arslan et al 2011**). However, our attempts to isolate these viruses using limiting dilutions and more passages did not result in reliable purified samples, making it impossible to proceed with further characterization.

**Figure 1:**
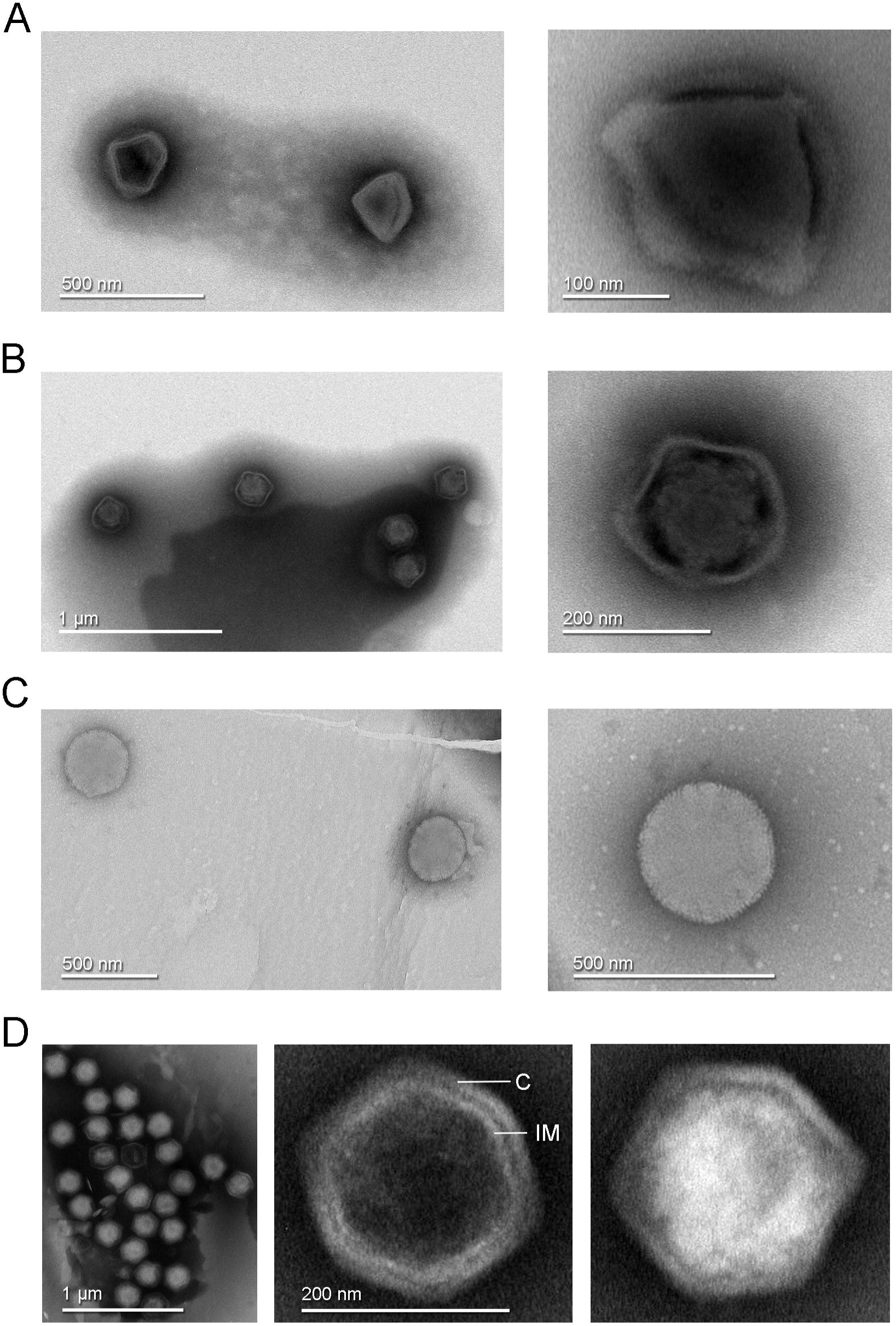
Transmission electron microscopy images of negative stained virus-like particles isolated from Finnish samples. A) Left, diamond-shaped virions found in a Recirculating Aquaculture System (RAS) tank sample and in experimental aquarium samples. Right, enlarged view showing the virion interior with at least three attachment points to the external capsid. B) Left, virions with loosely structured capsids found in a RAS tank sample and in experimental aquarium samples. Right, detailed view of one virion of this morphotype. C) Left, round morphotype found in an experimental aquarium sample; right, enlarged view of a spherical-shaped virion. D) Left, clusters of full and empty Jyvaskylavirus virions isolated from a composting soil sample. Center, an empty capsid showing the double-layered architecture of Jyvaskylavirus; C marks the capsid, and IM marks the putative internal membrane. Right, a fully packaged Jyvaskylavirus virion.

The isolate from sample 85 had a single morphotype of approximately 200 nm in size, displaying an icosahedral shape with particles that were mostly genome-filled and a few that were empty (**Figure 1D**). The empty particles possibly displayed an internal membrane beneath the capsid (**Figure 1D, centre**).

This sample was collected from the municipal Waste Treatment Centre, Jyväskylä, Finland. The sample was in soil state and originated from a mixture of 70% garden waste, 15% woodchips and 15% pre-treated biowaste. Virus Ac-85 was chosen as a model to represent the first giant virus isolate from Finland and named Jyvaskylavirus as a homage to the city of its isolation. Jyvaskylavirus has a fast replication cycle in its *A. castellanii* host, only causes CPE in *A. polyphaga* or *V. vermiformis* in high concentrations likely due to cytotoxic effect of the virus preparation, it is sensitive to chloroform treatment (10% for ten minutes) and is stable for long periods (up to 109 days) even at 37 °C (**Supplementary Material and Supplementary Figure 2)**. A fast replication cycle is a feature also shown for other marseilleviruses (**Boyer et al 2009; Fabre et al 2017**).

### Jyvaskylavirus belongs to the *Marseilleviridae* family

Jyvaskylavirus contains a 359,967 base pairs (bp) genome, with a 42.80% GC content and 388 predicted open reading frames (ORFs) coding for proteins with sizes that vary between 99 and 1,525 amino acids. The positive DNA strand codes for 186 ORFs whereas the other 202 are in the negative strand (**Figure 2A; Supplementary Table 2**). During genome annotation, some of the *Nucleocytoviricota* conserved proteins were detected, including the DNA polymerase family B (ORF177) and the A32-like packaging ATPase (ORF23). The most of Jyvaskylavirus genes (about 67%) code for uncharacterized proteins. The second major function category is the DNA replication, recombination and repair genes, including three histone-like proteins (ORFs 215, 216, 320), a typical marseillevirus genome feature (**Figure 2B**) (**Bryson et al 2022**). Three new ORFans were detected (ORFs 264, 265, 289), representing genes that have no significant similarity with any other sequences from the database used in this analysis. ORF 264, 265, and 289 codes for putative proteins with 108, 157 and 119 amino acids, respectively. We searched for translation-related genes, and found three translation factors, including a translation initiation factor (ORF 154), an elongation factor (ORF 318) and a peptide chain release factor (ORF 28). No tRNAs or aminoacyl tRNA synthetases genes were found. Furthermore, most of the BLASTp best hits for Jyvaskylavirus amino acid sequences matched Lausannevirus or Port-miou virus that are phylogenetically related to marseilleiviruses from lineage B. This observation can also be reinforced by the phylogeny based on DNA polymerase family B, which clusters the Jyvaskylavirus within *Marseilleviridae* family, together with other marseilleviruses from clade B (**Figure 2C, Supplementary Figure 3A**). When analyzing the genome synteny of different marseilleviruses genomes, it is shown that Jyvaskylavirus presents similarity blocks comparable to those from clade B marseilleviruses (**Supplementary Figure 3B**). Searching the Jyvaskylavirus major capsid protein and DNA polymerase sequences in the MGnify-database (**Richardson et al 2023**) yields multiple hits with significantly low E-values (< 1e-80), as expected from the apparent ubiquity of marseilleviruses. Of note was the detection of similar sequences in metagenomes and transcriptomes obtained from drinking water distribution systems of ground and surface waterworks in Central and Eastern Finland, evidencing that marseilleviruses are prevalent but still unexplored in this region (**Tiwari et al 2022**).

**Figure 2:**
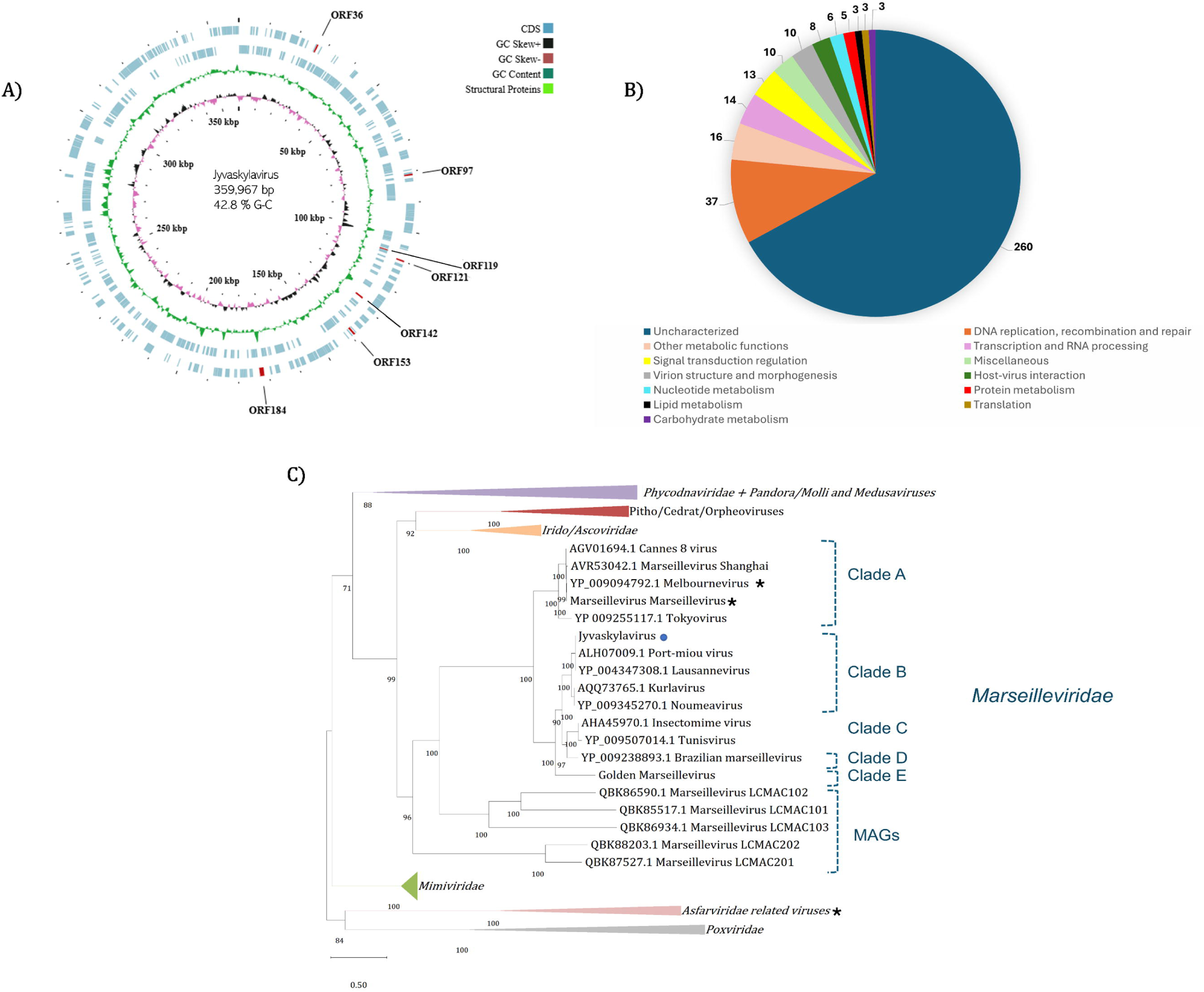
Jyvaskylavirus genomic data. A) Representative map of Jyvaskylavirus genome features. The G-C content, G-C skew, and ORFs distribution throughout the DNA sequence are coded by different ring colors as indicated in the color legend above. ORFs coding for the structural proteins mentioned in this paper are indicated by their ORF number. The outer blue ring represents the forward strand (positive sense) whereas the inner blue ring represents the reverse strand (negative sense). This illustrative genome map was constructed using CGview server (**Grant et al 2008**). B) Number of Jyvaskylavirus proteins according to the function predicted during genome annotation; n= 388. C) Maximum-likelihood phylogenetic tree based on DNA polymerase family B amino acid sequences from different nucleocytoviruses. The Jyvaskylavirus sequence is indicated by a blue circle. An asterisk (*) marks close viruses with structures obtained by cryo-EM. The alignment was performed with MUSCLE and the maximum-likelihood tree was reconstructed using IQtree software using ultrafast bootstrap (1000 replicates). The best-fit model selected using ModelFinder (implemented in IQtree) was VT+F+R5. Scale bar indicates the number of substitutions per site.

### Jyvaskylavirus attachment to host cells and extracellular virion clusters by scanning helium ion microscopy

To visualize the interaction of Jyvaskylavirus virions with its *A. castellanii* host cells we used a scanning Helium Ion Microscope (HIM). This imaging technology has proven successful in the study of bacteriophages and their host bacteria in the past (**Leppanen et al 2018**). We adapted the sample preparation methods to image cultures of *A. castellanii* infected with Jyvaskylavirus. Sample preparation was made by allowing cells colonize silicon chips treated with poly-l-lysine followed by infection and fixation, avoiding any need for cell scraping and pelleting for successfully imaging the amoeba cells and Jyvaskylavirus virions (**Figure 3**).

**Figure 3:**
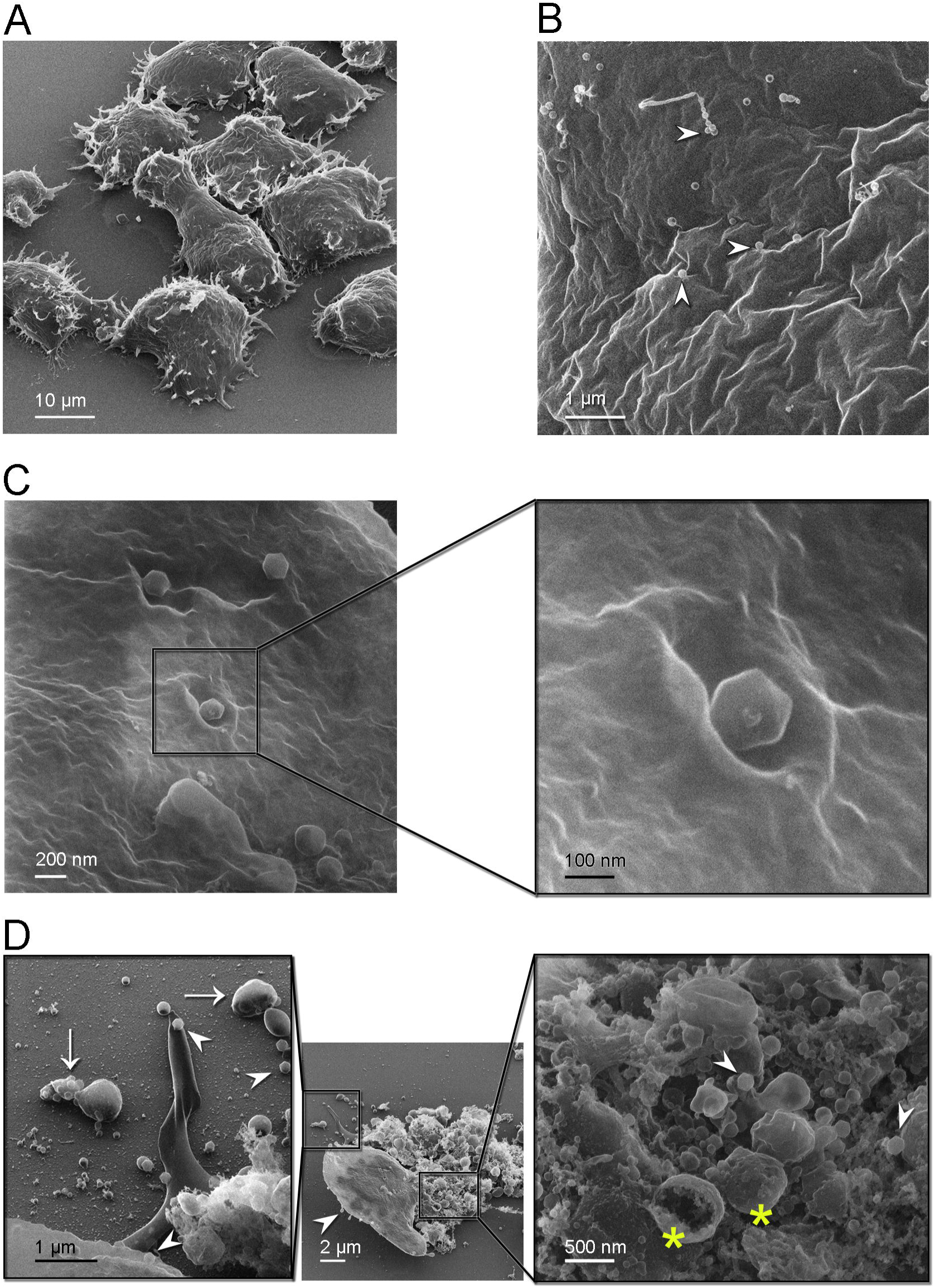
Helium ion microscopy images of Jyvaskylavirus attachment to *A. castellanii* cells. A) *A. castellanii* cells with spined structures (acanthopodia); elongated cell at the center. B) Details of a cell containing several viral particles on its surface (white arrowheads mark virions). C) Icosahedrally shaped virions near cell surface invaginations appearing as craters; inset, details of a virion likely starting the infection process through endocytosis. D) Centre, one cell containing viruses on its surface (white arrowhead) near a burst cell displayed with its ruptured content. Right inset, details of the burst cell content, showing several vesicles (yellow asterisks) and viruses (white arrowheads). Left inset, clusters of virions inside extracellular vesicles indicated by white arrows and individual virions by white arrowheads.

By HIM imaging *A. castellanii* cells appear as mainly oval-shaped with spined structures (acanthopodia) protruding from the cell surface; furthermore, a cell with an elongated shape was captured likely in the process of cell division (**Figure 3A**). No cysts were seen in these samples. At a closer range, cell surfaces appear rugose, full of crevices, and contains virions attached to it (**Figure 3B**). One of strategies for marseillevirus entry is the triggering of an endosomal-stimulated pathway (**Arantes et al 2016**). We captured a cell with virions attached in regions showing invaginations of the cell membrane, probably indicating the process of endocytosis (**Figure 3C**). These craters, surrounded by walls with varying degrees of steepness, differ in size although the estimated diameter to accommodate at least one virion is about 300 nm (**Figure 3C, inset)**. Alongside intact cells with attached virions, we also observed a broken cell nearby, with most of its cell contents released (**Figure 3D, centre**). Extracellular vesicles larger than 500 nm in size containing multiple virions inside were visualized close to burst cells (**Figure 3D, left inset**). These virion clusters are important for marseilleviruses, which are smaller than other giant viruses, to reach the required threshold for triggering amoebal phagocytosis and thus initiating their infection cycle through this entry mechanism (**Arantes et al 2016**). Details of the ruptured cell contents, revealing the presence of large internal spherical vesicles and intracellular virions, are shown in **Figure 3D** (right inset). Single virions were also observed on the substrate, suggesting that they could attach to foraging cells.

### Jyvaskylavirus forms intracellular vesicles and viral factories inside *A. castellanii* host cells

Intracellular details of the replication cycle of Jyvaskylavirus were imaged by transmission electron microscopy using ultrathin sectioning of cells undergoing cytopathic effect at 24 hours post-infection (hpi). At 24 hpi, the cells are overtaken by virus production, with intracellular vesicles of varying dimensions (*e.g* 1-2 µm), which contain already apparent virions (**Figure 4A**). Large viral factories are observed, with some occupying almost the entire cell area (**Figure 4B**). The nucleus and other cellular components, such as mitochondria, can be clearly distinguished within the cell sections (**Figure 4A-C**). Several individual virions also populated the cytoplasm. One imaged vesicle is juxtaposed to the nuclear membrane and contains several assembled virions with clear icosahedral shape. These virion-rich intracellular vesicles are likely the source of the extracellular ones imaged by HIM in **Figure 3D (left inset)**. Membrane-related structures located near or nearly attached to the luminal side of the large vesicle are discernible (**Figure 4C-D**). These membrane-derived assembled structures might serve during virus morphogenesis (**Figure 4D**). Inspection of the interior of viral factories unravels distinct stages of particle formation (**Figure 4E**). A putative assembly path can be extrapolated from the analysis of images of particles, which transition from a half-assembled icosahedron to a particle with an open vertex through the orderly accumulation of capsid proteins (**Figure 4F**). This open vertex is potentially used for genome packaging, and it is subsequently closed by the plugging of peripentonal and penton proteins.

**Figure 4:**
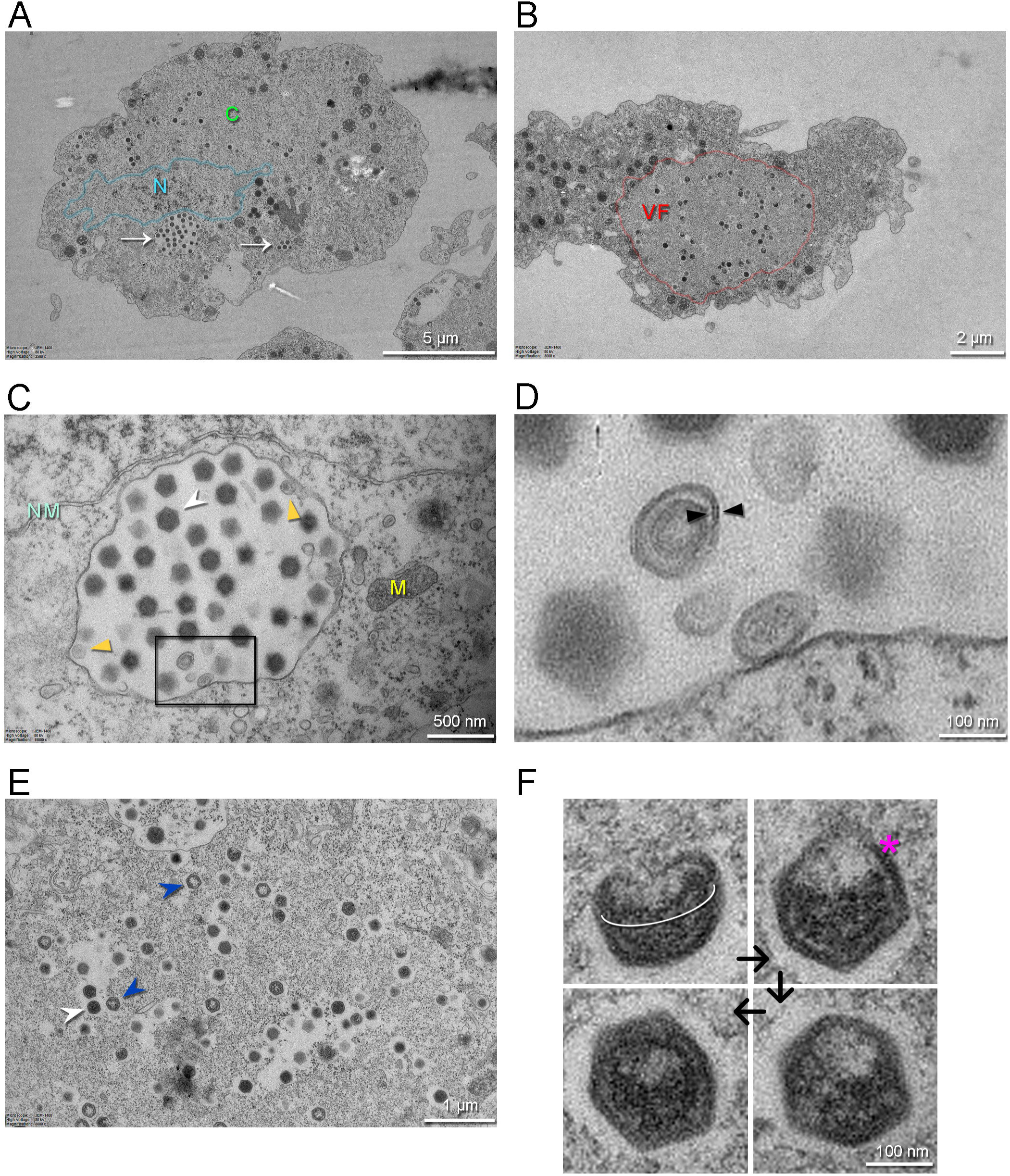
Transmission electron microscopy images of thin sections of *A. castellanii* cells infected by Jyvaskylavirus. A) Infected cell containing viruses spread over its cytoplasm marked by C (green) and with intracellular vesicles filled with viruses indicated by white arrows. The nucleus, whose boundary is highlighted in semi-transparent cyan, is indicated by the letter N (cyan). B) One infected cell with a large viral factory (VF, red) in its cytoplasm. C) View of an intracellular vesicle with icosahedral genome-filled virions as marked by a white arrowhead; membrane related structures nearby the vesicle interior are marked by dark-yellow triangles. NM (light cyan) and M (yellow) mark the nuclear membrane and the mitochondria, respectively. D) Enlarged view of the region marked by the black rectangle in C) showing possible membrane-related structure juxtaposed to or detached from the vesicle interior; black triangles possibly indicated a forming membrane vesicle. E) Details of virions in different stages of maturation inside the viral factory; DNA-full particles in white arrowheads, empty particles in blue arrowheads. F) Putative stages of virion assembly (indicated by the black arrows) as derived from the inspection of distinct particles in E and other cellular sections. The white elliptical line highlights a capsid aperture, while the red asterisk indicates an assembling capsid; in the remaining virion images, the capsid appears more assembled.

### Jyvaskylavirus three-dimensional architecture

Jyvaskylavirus icosahedral virion, determined to 6.3 Å resolution as judged by the gold-standard Fourier shell correlation possesses a diameter of ∼ 2,516 Å (vertex-to-vertex) and a triangulation number *T* = 309 (*h* = 7, *k* = 13) (**Figure 5A**, **Supplementary Figure 4 and Supplementary Table 3**). The protein capsid shell, approximately 120 □ thick, can be geometrically represented by trisymmetrons and pentasymmetrons as similarly done with other NCLDVs (**Figure 5B**) (**Sinkovits & Baker 2010**). The trisymmetron and pentasymmetron comprises 136 and 30 pseudo-hexameric capsomers, respectively, along with one penton complex, containing five copies of the penton protein (**Figure 5B-C**). Jyvaskylavirus icosahedral asymmetric unit (IAU) is composed of 51 pseudo-hexameric capsomers plus 1/3 of the capsomer sitting on the icosahedral 3-fold axis (**Figure 5C**). Both the capsid organization and virion size are similar to those of other marseilleviruses, such as Melbournevirus and Tokyovirus. Pacmanvirus, considered to be at the crossroads between *Asfarviridae* and Faustoviruses, also possesses the same *T* number (309) and a comparable diameter to Jyvaskylavirus. In contrast, other giant viruses, such as African swine fever virus (ASFV), representative of the *Asfarviridae* family, have a *T* number of 277 and a diameter of approximately 2,100 Å, while PBCV-1, a member of the *Phycodnaviridae* family, has a *T* number of 169 and an average diameter of 1,900 Å. All of the above-mentioned viruses have been shown to possess a major capsid protein with a vertical double jellyroll (DJR) fold that composes the capsid shell, along with an internal membrane bilayer. Minor capsid proteins have been identified and structurally modelled for the smaller virions ASFV and PBCV-1 (Wang et al. 2019; Shao et al. 2022).

**Figure 5:**
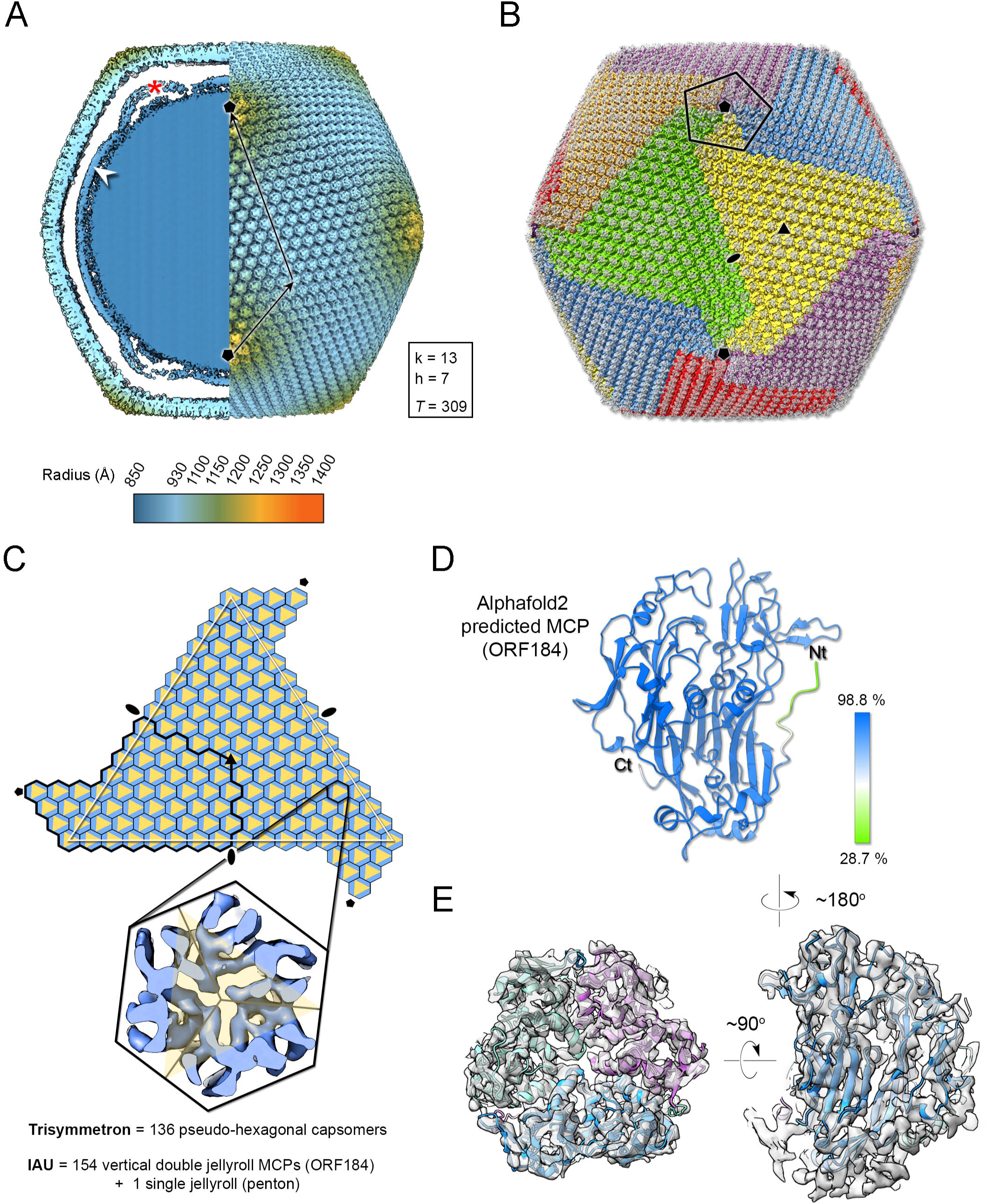
Jyvaskylavirus cryo-EM reconstruction. A) Left, a central slab of the isosurface of the 3D density map, downsampled to a pixel size of 5.36 Å, low-pass filtered to 15 Å, and normalised, showing the interior of the virion colour-coded by radius as from key. The white arrowhead marks the membrane bilayer, while the red asterisk highlights the membrane bulging beneath the five-fold vertex. Right, isosurface of half the virion with black arrows indicating the triangulation indices h=7, k=13 (*T* = 309). A threshold level of 0.6 was used in ChimeraX to render both views. B) Representation of the virion using trisymmetron geometry, with each trisymmetron color-coded differently, and a pentasymmetron marked by a pentagonal black line. C) A schematic of a virus facet with a trisymmetron marked with a white triangle with three icosahedral asymmetric units (IAU), one of which is marked by a thick black line. The pseudo-hexameric morphology displayed by a capsomer is represented by a hexagon coloured in light blue, while the true trimeric state of the MCP is depicted as a yellow triangle. To build the IAU (excluding the penton protein), 51 pseudo-hexameric capsomers and one-third of the capsomer located at the 3-fold symmetry axis are required, resulting in a total of 154 major capsid proteins (MCP) forming the IAU. The inset shows a cut-through of the density along the three-fold axis of a capsomer. D) Alphafold prediction of the MCP ORF184 shows, with very high confidence, that the fold adopted by the ORF is a vertical double jellyroll. E) Trimeric model of the capsomers rigid-body fitted into the original cryo-EM density and rendered in ChimeraX; left, viewed along the trimer fold axis and on the right, viewed orthogonally to it. The three copies of the MCP are represented in cartoon and coloured in green, light magenta, and light blue, while the corresponding density is shown in white transparent surface.

Beneath the Jyvaskylavirus capsid, the membrane vesicle follows icosahedral symmetry, although it also displays a high degree of sphericity (**Figure 5A, left**). This internal membrane vesicle encloses the genome. Some particles, excluded during 2D classification, showed heterogeneous membrane morphologies indicating their structural fragility. At the five-fold axis between the capsid and the membrane, there is relatively weak density suggesting the presence of additional proteins (see below), along with a clear bulging of the membrane vesicle at the five-fold (radius of curvature of ∼275 Å) (**Figure 5A, left**). To the best of our knowledge, this bulging has only been clearly observed in the closely related Melbournevirus and Tokyovirus.

### Jyvaskylavirus structural proteins composing the capsid

At 6.3 Å resolution alpha-helical secondary structural motifs are identifiable while the separation of β-strands became clearer beyond 5 Å. Our density corresponding to the capsomer displays unequivocally pseudo-hexameric morphology with the characteristic footprint of trimers formed by vertical double jellyroll seen in other viruses of the kingdom *Bamfordviriae* (**Andres et al 2020, Ravantti et al 2020, Simmonds et al 2023**). A cross-section of the density shows the β-barrel walls, further supporting that the major capsid protein (MCP) possesses a vertical DJR fold (**Figure 5C, inset**). During genome annotation based on sequence homology with other marseilleviruses, ORF184 was identified as a potential MCP. We submitted ORF184 to AlphaFold3, which predicted a model with a DJR fold (**Figure 5D**) (**Abramson et al 2024**). The fitting of this model into density leaves no doubts about ORF184 being the MCP and a total of 154 copies of the MCP compose the IAU (**Figure 5C, E**).

Then, we structurally identified the penton protein of Jyvaskylavirus as ORF142 by using the latest version of ModelAngelo software with the hidden Markov model (HMM) search procedure against the publicly available cryo-EM block-reconstructed capsid densities of the Melbournevirus, a clade A *Marseilleviridae*, at ∼3.5 Å resolution (EMD-37188, 37189, 37190) (for details see Materials and Methods and **Supplementary Figure 5**) (**Jamali et al 2024**). This identification was based on the hypothesis of structural conservation among conserved capsid components and partial protein sequence conservation among members of the *Marseilleviridae.* Model prediction of the ORF142 in Alphafold3 produced a compact eight stranded β-barrel (BIDG – CHEF) typical of a jellyroll, elaborated by extended loops between strands DC and EF and a long C terminal tail (**Figure 6A, left**). The jelly-roll core of the penton proteins showed a reasonable fit into the corresponding Jyvaskylavirus densities at 6.3 Å resolution (**Figures 5E and 6A**).

**Figure 6:**
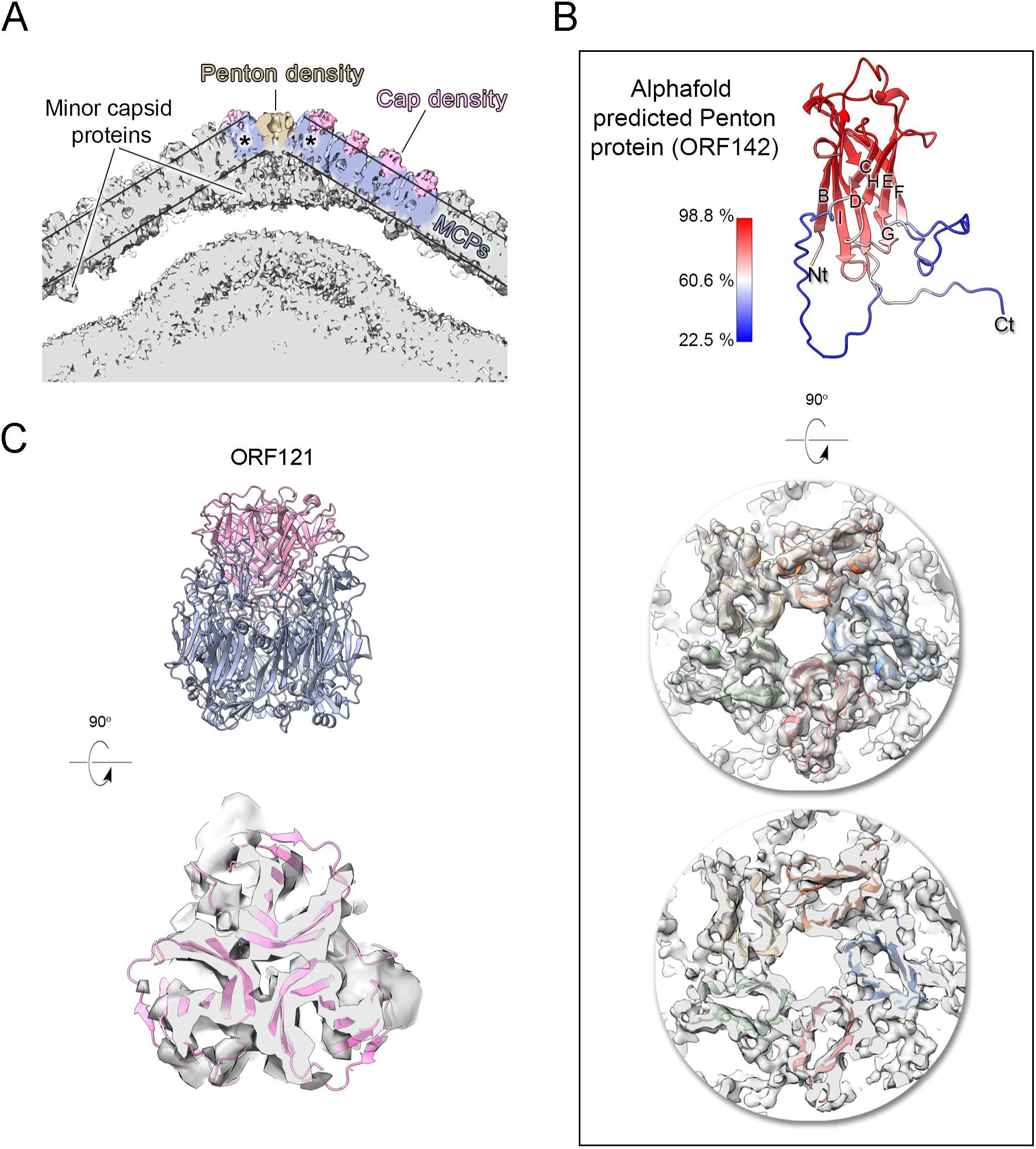
Jyvaskylavirus penton and capsomer-cap proteins. A) Cut-through view of the virion cryo-EM density (binned twice from original size) at the 5-fold axis rendered in grey in ChimeraX, with some regions of the capsid shell coloured to highlight the penton density (light brown), the major capsid protein densities (slate-blue) and within the parallel outlined black lines and the cap densities (pink), other regions beneath the MCPs and penton correspond to different minor capsid proteins; the black asterisks mark the peripentonal capsomers. B) Top, a cartoon representation of the predicted model of the penton protein, with strands labeled BIDG/CHEF and color-coded by confidence level. Center and bottom, penton complex fitted into the density, shown from the top and as a cut-through, respectively. C) Top, atomic model of the trimeric cap in pink cartoon, composed of three copies of ORF121 with a β-barrel fold inserted on top of the pseudo-hexameric capsomer, shown in slate-blue cartoon. Bottom, cut-through view of the cap model fitted into the original density, Gaussian filtered and rendered as semi-transparent grey in ChimeraX.

Additional density was observed atop the pseudo-hexameric capsomers, capping the central region formed by the ORF184 jellyroll towers, and likely stabilizing the trimer from the exterior. Using the same methodological strategy as for the penton protein, we identified ORF121 as this cap protein, which possesses a β-barrel fold and forms a trimer, with its N-terminal ends inserting into the crevice formed by the MCP jellyroll towers (**Figure 6B**). Beneath the capsid shell, weaker density is visible at varying distances from the virus center and at different locations within the trisymmetron (**Figures 6B and 7A, top left**). Their positioning resembles that seen in Tokyovirus and Melbournevirus, which have been linked to pentasymmetron protein components (PCs) and minor capsid proteins (mCPs) with proposed roles in scaffolding, cementing, and zippering (**Chihara et al 2022; Burton-Smith et al 2021**). However, no information is currently available regarding their corresponding ORFs and/or 3D models.

**Figure 7:**
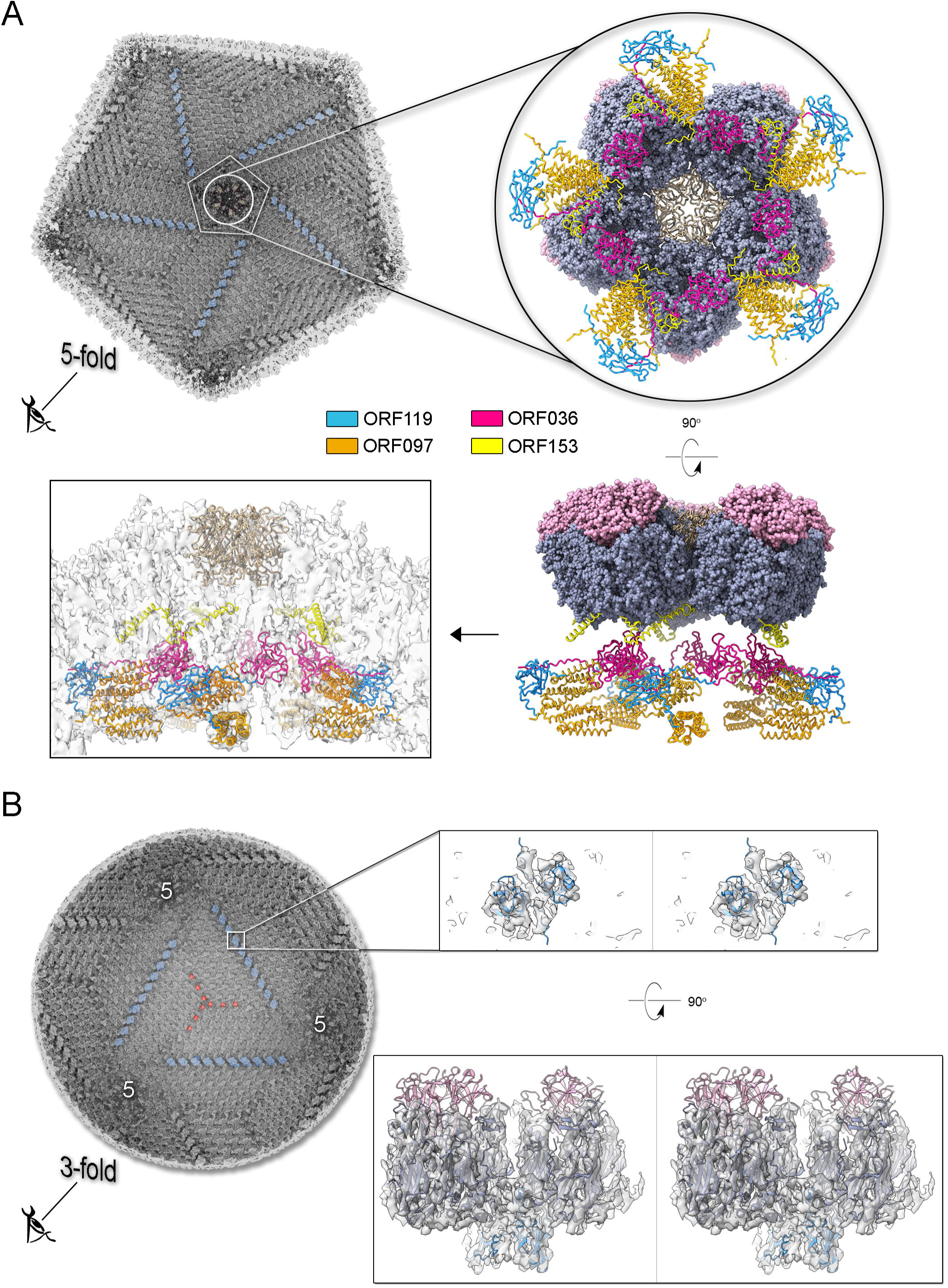
Jyvaskylavirus pentasymmetron protein components and trisymmetron facet glueing protein. A) Top left, view of the cryo-EM density of Jyvaskylavirus (binned four times from the original size and Gaussian filtered) as seen from within the virion along the five-fold icosahedral symmetry axis. The white pentameric line marks the pentasymmetron region from below, while the white circle highlights the density corresponding to some identified proteins. Further densities corresponding to cementing, zippering and lattice scaffolding proteins are also visible, with one of them coloured in dodger-blue (see also panel B). The enlarged inset (top right and below) shows the spatial organization of the four identified ORFs and modelled using AlphaFold3, represented as cartoon tubes and coloured as per the legend. The penton proteins are coloured light brown, while peripentonal capsomers and capsomer-cap proteins are shown as space-filled atoms and coloured as slate-blue and pink, respectively. Left bottom, the different protein components fitted into the original 6.3 Å resolution Jyvaskylavirus cryo-EM density (white semi-transparent) binned to 2.68 Å/pix and rendered in ChimeraX; MCPs and capsomer-cap proteins have been omitted for clarity (for fitting metric see Materials and Methods). B) Left, view of the cryo-EM density of Jyvaskylavirus, as shown in the top left panel of A), but along the three-fold icosahedral symmetry axis. A cementing protein, colored red, runs through the capsomers; however, the corresponding ORF has not been identified. Densities colored dodger-blue, which glue the capsomers across two trisymmetrons, correspond to ORF119, as shown in the large inset on the right. In this inset, stereo view of dimers of ORF119 are depicted as dodger-blue cartoons fitted into the original cryo-EM density, Gaussian filtered and rendered as semi-transparent grey in ChimeraX. At the bottom right, a 90-degree stereo view shows the spatial arrangement between two capsomers (navy blue), with the cap protein ORF121 (pink cartoon) positioned on top, and ORF119 fitted into the density at the base of the adjacent MCPs.

Using the above-mentioned approach (**Supplementary Figure 5**), we identified four additional Jyvaskylavirus proteins - ORF36, ORF97, ORF153, and ORF119 – located beneath the pentasymmetron capsomers. We predicted their 3D models using AlphaFold3 and positioned them within the corresponding higher-resolution Melbournevirus densities, where the core of the different molecules generally fit well within the constraints of the map (**Supplementary Figures 6-8**). We then placed all the newly identified and predicted Jyvaskylavirus 3D models into our cryo-EM density map at 6.3 Å resolution (**Figure 7**). Intriguingly, ORF119 was also identified as the protein that runs along the edge of the trisymmetron facets, acting as a glue between adjacent capsomers belonging to two different trisymmetrons (**Figure 7B** and **Supplementary Figure 8**). The relative orientation of the ORF119 molecule beneath the pentasymmetron and along the edges of two trisymmetrons is about 90 degrees. However, not all the visible density beneath the capsid in the deposited Melbournevirus block-based reconstructed maps could be accounted for by the fitted structures. Interestingly, all Alphafold3 predicted ORFs presented unordered regions, mostly at the N- and C-terminal ends, with lengths varying depending on the protein and their location. While we did not attempt further remodelling of these flexible regions into Melbournevirus densities, a more complete representation of Melbournevirus is achievable. To this end, we identified the corresponding Jyvaskylavirus ORFs in Melbournevirus through sequence comparison with Melbournevirus isolate 1 (NCBI Reference Sequence: NC_025412.1) (**Supplementary Table 4**). However, when the identified Jyvaskylavirus ORF sequences were analyzed using BLASTp without restricting the search to the Melbournevirus reference, many hits were observed in other giant viruses, primarily marseillevirus. Remarkably, some of these hits scored higher than those for Melbournevirus, supporting the presence of homologous proteins in these viruses (**Supplementary Table 5**).Further, a distinctive feature of the internal membrane vesicle is the bulging of the lipid bilayer at the five-fold equivalent to that found in Tokyovirus (**Figure 5A, left** and **Figure 6A**) (**Chihara et al 2022**). The identified PCs crown this region, and it is plausible that, along with yet unidentified proteins, they act as effectors of the bulging by tethering the membrane. The radius of the bulging is likely related to the extent of their localization beneath the pentasymmetron.

## DISCUSSION

Jyvaskylavirus is the first characterized giant virus from Finland. So far the only other isolated giant viruses from the Nordic Countries are the still uncharacterized Lurbovirus from Sweden and a collection of viruses capable of infecting microalgae isolated in Southern Norway (**Kördel et al 2021**; **Sandaa et al 2001; Johannessen et al 2015**). There is genomic evidence of NCLDV presence in the Greenland ice sheet and of a high diversity of giant viruses, including the detection of marseilleviruses, in the Loki’s Castle deep-sea vent located in the Mid-Atlantic Ridge between Iceland and Svalbard (**Perini et al 2024**; **Backstrom et al 2021**). However, no virus was isolated in these studies. Jyvaskylavirus belongs to clade B of the *Marseilleviridae* family, so far making it the northernmost known member of the family. The first marseillevirus was isolated from France in 2007 (**Boyer et al 2009**). Now there are more than 60 isolates known, obtained from varied sample sources in seven countries over five continents (**Sahmi-Bounsiar et al 2021**). Our genetic analysis showed that the Loki’s Castle marseilleviruses group together and that they are not close to Jyvaskylavirus, probably forming unique clusters among themselves (**Supplementary Figure 3**). Jyvaskylavirus is also unique for being the first found from a composting sample, demonstrating that these giant viruses are found in water and soil samples worldwide.

For the characterization of Jyvaskylavirus we integrated complementary imaging techniques, including the use of Helium Ion Microscopy for imaging the virus and its *A. castellanii* host. This demonstrates the applicability of this technique to giant viruses and sets the methods for sample preparation, opening new ways for the study of giant viruses and interactions with their host.

Ultrathin TEM sectioning enabled us to capture snapshots of a putative assembly pathway, unravelling the progressive formation of the capsid. Single-particle cryo-EM provided critical insights into the structure of Jyvaskylavirus and its evolutionary relationship with other viruses. The 3D reconstructed cryo-EM density at 6.3 Å resolution clearly recapitulates the architecture (*T* = 309; vertex-to-vertex diameter 2,500 Å) and capsid protein organization observed in Melbournevirus and Tokyovirus, other members of the *Marseilleviridae* family (**Burton-Smith et al 2021; Chihara et al 2022**). Although it is not possible to distinguish individual β-strands at this resolution, with the aid of Alphafold3, we unequivocally identified and placed the predicted ORF184 MCP vertical double jelly-roll fold model into density. Additionally, we identified and predicted the fold of the penton protein encoded by ORF142 to adopt a vertical jellyroll topology. This model nicely fitted the corresponding density, plugging the center of the pentasymmetrons. There is structural conservation of the penton protein fold with that of other viruses such as penton proteins P31 in bacteriophage PRD1 or VP9 in archaeal virus HCIV-1 of the *Bamfordviria* and *Helvetiavirae* kingdoms, possessing vertical double and single jellyroll MCPs, respectively (**Abrescia et al 2004**; **Santos-Perez et al 2019**). However, in ORF142, the CHEF strands are predicted to be tilted relative to the BIDG strands, with an estimated angle of approximately 60° based on visual inspection (**Supplementary Figure 9**). This type of penton complex acts like a plug-and-play system, able to incorporate various host-recognizing vertex proteins. These proteins are interchangeable and adapt to environmental evolutionary pressures (**Gil-Carton et al 2015**). The clear presence of a network of scaffolding, cementing, and minor proteins beneath, previously observed in ASFV and Tokyovirus, but also in smaller viruses such as PBCV-1, adenovirus, and PRD1, reiterates the universal requirement for the assembly of larger virus of an increase number of ancillary proteins tethering the MCPs at the vertices, within the facets and along the facets **(Abrescia et al 2004, Burton-Smith et al 2021; Liu et al 2010, Wang et al 2019, Chihara et al 2022, Shao et al 2022**). In this study, through cross-structural and sequence comparisons with available Melbournevirus block-based reconstructed densities and combination of AI-based modelling software (**Zhu et al 2018**; **Burton-Smith et al 2021**; **Jamali et al 2024**), we were able to elucidate the identity and the fold of five additional viral capsid components in Jyvaskylavirus: ORF36, 97, 119, 153 and 121 (and the corresponding ORFs in Melbournevirus). The former four are located beneath the pentasymmetron while ORF121 is the protein capping on the top the capsomers and it shares its fold with marseillevirus Noumeavirus NMV_189 protein (PDB ID 7QRR) (*e.g.*, rmsd 1.5 Å, 146 Cα aligned) and virophage Zamilion vitis protein Zav19 (PDB ID 7QRJ).

Remarkably, two ORF119 molecules that form a homodimer are also used to glue at their bases capsomers located at the edge of adjacent trisymmetrons. The orientation of the ORF119 molecule below the pentasymmetron differs from that along the edge of the trisymmetron, indicating that its binding mode depends on the structural environment. The reuse of the same protein in different regions of the facet for multiple functions highlights ORFs optimization and genetic parsimony. Particularly intriguing is that while AlphaFold3 predicts the core of most identified proteins, satisfying the constraints of the map, some regions are poorly ordered. One clear case is ORF153 which is predicted to possess about 40% of the residues unstructured. We propose that these regions together with flexible terminal ends are not merely a limitation of AlphaFold3’s predictive capabilities but rather reflect specific functional characteristics of these proteins, which may fold and adapt through protein partnering. Additionally, the observation that the identified Jyvaskylavirus minor capsid protein sequences are shared across other marseillaviruses supports their essential structural and stabilizing roles in these viruses.It has been shown that the conserved flexibility and function of intrinsically disordered proteins despite their often-fast rate and tolerance for mutations, plays critical physiological roles across all organisms and viruses (**Brown et al 2011**). Protein P30 of the lipid-containing bacteriophage PRD1, structurally determined more than 20 years ago, and predicted as a disordered protein, was the first to be identified possessing a string-like structure running through adjacent virus facets, interacting with capsomers along the edges and cementing them together (**Abrescia et al., 2004**). Giant viruses, however, are structurally more complex, with a larger number of minor capsid proteins holding the virion together beneath the pentasymmetron and trisymmetron. While our current study advances our understanding of this complexity by identifying various ORFs and positioning their models in the density map, deciphering their atomic interactions and specific roles in particle assembly will require higher resolution for the different components.

Jyvaskylavirus isolation is the first step in better understanding the giant virus diversity in the boreal environment. Our preliminary bioprospection attempt using a locally isolated amoebal host suggested a high diversity of unique viral like particles, while our screening of fish tank samples revealed mixed populations of other novel viruses with unique morphologies. The origin of the viruses found in aquarium water is unknown, and they could be originated from the ground water used to keep the fish or the fish microbiome itself. marseillevirus DNA polymerase sequences are present in metagenomes from Finnish drinking water distribution systems (**Tiwari et al 2022**) hinting to a wide distribution of these viruses and still unknown ecological role in Central and Eastern Finland. In addition, electron microscopy analyses revealed high structural diversity of giant viruses and virus-like particles in soil, suggesting a gap in knowledge in the diversity and ecology of environmental virus-host-interactions (**Hylling et al 2020**; **Fischer et al 2023**). Despite the technical challenges that prevented us from characterizing all isolated viruses in this study, our findings underscore the importance that broader sampling and new isolation attempts are essential to determine the distribution of these viruses. The use of host species isolated locally might avoid isolation bias towards the reference strains giving a better idea of viral diversity and allowing the description of unique viral groups.

In summary, marseilleviruses found in diverse locations and environments worldwide show a strictly conserved architecture, using equivalent pseudo-hexameric building blocks and a plethora of ancillary proteins. We suggest that this adaptation relies on the number, fold, and length of disordered regions in the minor capsid proteins, as well as the membrane composition. These factors not only facilitate correct assembly but would also modulate the virus’s structural stability across various environments.

## MATERIALS AND METHODS

### Key Resources Table

| Key Resources Table |  |  |  |  |
| --- | --- | --- | --- | --- |
| Reagent type (species) or resource | Designation | Source or reference | Identifiers | Additional information |
| Strain<br>*Acanthamoeba castellanii* | *Acanthamoeba castellanii* | Aix Marseille University, France | AC | Donated by Professor Bernard La Scola (Aix Marseille University, France). |
| Strain<br>*Acanthamoeba polyphaga* | *Acanthamoeba polyphaga* | Aix Marseille University, France | AP | Donated by Professor Bernard La Scola (Aix Marseille University, France). |
| Strain<br>*Vermamoeba vermiformis* | *Vermamoeba vermiformis* | Aix Marseille University, France | AP | Donated by Professor Bernard La Scola (Aix Marseille University, France). |
| Strain<br>Jyvaskylavirus | Jyvaskylavirus | This study | AC85 | Finnish clade B<br>marseillevirus.<br>Genome<br>accession<br>number:<br>PQ284187 |

### Samples used for screening, amoebal host cultivation and viral isolation

To get an overview on presence of giant viruses in Finland and their virus-host interactions, water and soil samples were collected from Jyvaskyla (Finland) and used fresh for viral isolation. Previously collected and stored samples were also used, e.g. aquaculture water samples, recirculating aquaculture filter pellets and frozen water from experimental aquaria (**Runtuvuori-Salmela et al 2020**; **Almeida et al 2019a**; **Almeida et al 2019b**). Sample details can be found at the **Supplementary Table 1**. Water samples were collected directly into Eppendorf tubes. Solid samples like soil and composting material (roughly one fourth of an Eppendorf tube) were resuspended in 1ml of PAS buffer (120 mg NaCl, 4 mg MgSO_4_.7H_2_O, 4 mg CaCl_2_.2H_2_O, 142 mg Na_2_HPO_4_, 136 mg KH_2_PO_4_ in 1 litre of water) and strongly vortexed for resuspension (**Thomas et al 2006**). Each sample had the following antibiotic mix added to avoid fungal and bacterial contamination during the isolation process: penicillin (0.14 mg/ml), gentamycin (50 mg/ml), amphotericin B (0.25 µg/ml), ciprofloxacin (0.004 mg/ml), vancomycin (0.004 mg/ml) and doxycycline (0.02 mg/ml). Indicated concentrations are the final concentration of each antibiotic in solution.

Isolation was made using *Acanthamoeba castellanii*, *A. polyphaga* and *Vermamoeba vermiformis* hosts. All three host strains were kindly donated by Dr. Bernard La Scola and grown using PYG media at room temperature (∼25 °C) (**Jensen et al 1970**). For the isolation process, cells were mixed with 50 µl of the samples in 96 well plates. Cell density was controlled by resuspending confluent T25 flasks in 1 milliliter of PYG media and diluting it 1/1000 before adding 200 µl to each well in the plates. The antibiotics mentioned above were also present in PYG media used for preparing the isolation plates. Cells were monitored daily for five days for the appearance of cytopathic effect (CPE). In case no effect appeared, the plates were frozen, thawed and 25 µl of each well used as samples for a second passage. Three passages of all samples were made in all three hosts. Negative samples were those that no CPE appeared after the third passage.

### Growth curve, chloroform sensitivity and stability

Growth curves were made by infecting confluent cells in 96 well plates with a low MOI (1/1000 dilution of the virus stock). For each dilution four wells were prepared. At the indicated time points the samples were frozen and then, after two freeze-thaw cycles, 50 µl of each well were transferred to a new 96 well plate containing confluent *A. castellanii* cells for titration. Chloroform sensitivity was made by exposing a 900 µl viral aliquot to 10% chloroform for ten minutes, inside an Eppendorf tube. As control, 900 µl of the same viral stock was mixed with 100 µl of PAS and incubated for ten minutes. After incubation both were serially diluted and titrated (four replicates per dilution). Stability was evaluated by diluting a viral stock serially (-1 to -11, 1ml aliquots) and storing the viral dilutions at room temperature (∼25 °C), cold room (∼8 °C) and inside an incubator (37 °C). Samples were stored for 109 days and then titrated. The incubation time used is this long because of the SARS CoV2-related lockdowns that happened during this experiment. All titers were calculated by tissue culture infectious dose (TCID)50 (**Reed and Muench 1938**).

### Sequencing and genomic data

For DNA extraction, a confluent T25 was infected at a low MOI and the supernatant was harvested after the appearance of full CPE. Two aliquots of 2ml each were subjected to nuclease treatment and capsid precipitation by ZnCl_2_ (**Santos 1991**). After proteinase K treatment the samples were mixed with ethanol:guanidine and DNA extraction was finished using the GeneJET Genomic DNA Purification Kit (Thermo Fisher). DNA paired-end libraries (2□×□250-bp) were constructed with 1□ng of the viral genome as input using the Nextera XT DNA kit (Illumina, Inc., San Diego, USA) and sequenced on the Illumina MiSeq for 39-hours, the same strategy employed by **Brahim Belhaouari et al 2022**.

The assembled genome was submitted to open reading frames prediction (ORFs) using GeneMarkS (**Besemer et al 2001**). Also, a search for tRNA genes was performed using ARAGORN program (**Laslett & Bjorn 2004**). Only ORFs having more than 150 nucleotides were considered to analysis. Similar sequences for each predicted protein were searched using BLASTp (expect threshold: 10-3) against NCBI non-redundant protein sequences (nr) database. To perform synteny analysis, complete genome sequences from different marseilleviruses were obtained in GenBank. The sequences underwent manual curation to correct distortions caused by the circular topology of marseilleviruses genomes. Then, synteny analysis were conducted using MAUVE program with default parameters (**Darling et al 2004**).

Phylogenetic trees were constructed based on DNA polymerase and major capsid protein (MCP) amino acid sequences. The sequences used for alignments were obtained using BLASTp (expect threshold: 10-3) against NCBI non-redundant protein sequences (nr) database. The alignments were performed using MUSCLE executed through MEGA X program (**Edgar 2004; Kumar et al 2018**). Maximum-likelihood phylogenetic trees were constructed using IQ-tree software (version 1.6.12) with 1,000 bootstrap replicates as branches support (**Nguyen et al 2014**). Best-fit substitution models were selected by ModelFinder algorithm implemented in IQtree (**Kalyaanamoorthy et al 2017**). The phylogenetic trees were visualized using iTOL (**Letunic & Bork 2021**).

### Microscopy

Samples with noticeable CPE during the viral isolation process were prepared for checking the presence of negatively stained viral-like particles (VLPs). Two microliters of the lysates (supernatant straight from the isolation plate) were added to a microscopy grid and incubated for two minutes at room temperature. The excess liquid was removed with a water soaked Whatman paper. Then five microliters of 2% phosphotungstic acid was added to the grid and its excess was removed after an incubation of two minutes. The grids were left for drying at room temperature for five minutes and used for imaging at a Jeol JEM-1400 electron microscope straight away.

Samples destined for thin sectioning were prepared by infecting a T75 flask with a low multiplicity of infection (MOI). Twenty-four hours later, when the cytopathic effect started to appear, cells were harvested from the flask and pelleted by centrifugation (10 minutes at 2000 g). The pellet was resuspended in 2.5 % glutaraldehyde 0.1 M sodium phosphate buffer and kept under slow rotation for one hour at room temperature. After the incubation the cells were pelleted again, resuspended in 0.1 M sodium phosphate buffer and sent for blocking and thin section preparation. The samples were also imaged at a Jeol JEM-1400 electron microscope.

Substrates for helium ion microscopy were prepared by incubating silicon chips in poly-l-lysine (Sigma Aldrich) for five minutes followed by two wash steps in sterile water and left to dry overnight. *A. castellanii* cells were seeded in 24 well plates containing the poly-L-lysine coated substrates and infected at different MOI. After the appearance of cytopathic effect, the samples were fixed with 2.5 % glutaraldehyde in 0.1 M sodium cacodylate for two hours followed by three washes wash with 0.1 M sodium cacodylate buffer. After that, the samples were stained by incubation for one hour with 1 % osmium tetroxide followed by three washes in 0.1 M sodium cacodylate and a second staining with 0.1 % tannic acid for twenty minutes. Dehydration was done with sequential exposure to increasing concentrations of ethanol, using the following percentages: 35, 50, 70, 85, 95 and 99 %. Each exposure was made for thirty minutes, except for 99 %, which had one thirty minutes exposure followed by a second overnight exposure to ensure proper dehydration. Ethanol washing was followed up by a critical point drying step using twenty-four cycles in a Leica EM CPD300 equipment. Macroscopic sample structure did not change during the drying. Imaging was made using the Zeiss Orion Nanofab Helium Ion Microscope from the University of Jyväskylä Nanoscience Center with acceleration voltage 30 kV and ion current 0.2-0.3 pA. Electron flood gun was used during imaging to mitigate positive charging by the ions.

### Virus production for cryo-electron microscopy and data collection

A large stock of Jyvaskylavirus was prepared and purified for cryo-EM analysis. Twelve confluent T75 flasks were infected at a low MOI. After the appearance of full CPE the flasks were frozen-thaw to lyse still intact cells, and then all the flask contents were moved to falcon tubes. After one additional freeze-thaw cycle, a brief centrifugation (500 g, 5 minutes, 10 °C) was made to clear the lysate from cell debris. Then the viruses were pelleted (10.000 g, 60 minutes, 10 °C) and resuspended in 700 µl of PAS. The resuspended viruses were loaded into a 10-50 % sucrose gradient and centrifuged (6500 g, 90 minutes, 10 °C). The viral band of the gradient was collected, mixed with PAS buffer to dilute the sucrose and the viruses were pelleted again. The final pellet was resuspended in 150 µl of PAS buffer, aliquoted and shipped at 10 °C to the CIC bioGUNE for cryo-EM analysis. Upon arrival the sample was vitrified either using a Vitrorobot Mark IV (Thermofisher) or an Automatic Plunge Freezer EM GP2 (Leica). As a ‘quality control’ step, some of the grids were inspected using the in-house JEM-2200FS/CR (JEOL, Ltd.) cryo-TEM equipped with a K2 bioquantum camera. The remaining grids were shipped for high-resolution imaging at eBIC - Diamond Light Source (Didcot, UK) in line with democratic access to large infrastructure (**Stuart et al 2016**). Four data collections on a Titan Krios 300 kV with a K3 camera were performed at a nominal magnification of ×64,000, resulting in a final pixel size of 1.34 Å/pix or 1.35 Å/pix depending on the microscope used (**Supplementary Table 3**). Briefly, samples were vitrified on Quantifoil Cu R2/2 or R2/1 300 mesh grids and then collected over 40-45 fractions (with a dose per frame of ≈ 1 e-/Å^2^), with defocus ranges from -0.6 to 3 µm. As we had approximately one particle per hole, different software were used at each data collection to test which strategy would yield the highest number of particles (EPU, TOMO5 and Serial EM; for details in each data collection see **Supplementary Table 3**). After four data collections we obtained 3,742 useful particles from 17,720 movies.

### 3D reconstruction of Jyvaskylavirus

MotionCorr2 was used to correct the induced beam-shift across the frames within the movies, while the individual movies’ CTF was estimated using the CTFFIND4 software (**Rohou & Grigorieff 2015; Zheng et al 2017**). Particle auto-picking for the first data collection was performed in crYOLO, training the model with 10 movies (**Wagner et al 2019**); however, the picking process was supervised. For the remaining acquisitions, particles were manually selected in RELION 3.1 due to the limited number of movies available (**Scheres 2011**). Virions were extracted into a 1,000 x 1,000 pixel box, resulting in a final pixel size of 3.087 Å. For each dataset, several rounds of 2D and 3D classifications were performed before 3D refinement, with the initial 3D reference being generated *ab-initio* from a limited number of particles from the first data collection. Subsequently, the particles were re-extracted and re-centered, then combined with those from the other data collection after undergoing the same pre-processing workflow. Further classifications led to a homogeneous class comprising 3,742 particles (**Supplementary Table 3**), which then underwent 3D icosahedral refinement with the original pixel size (1.34 Å/pix) on a 2,304 pix box. This was achieved by using the Picasso HPC at the University of Malaga node of the Spanish Supercomputing Network (RES). Computing resources used for the above refinement included 3 tasks, 32 cpu per task and 1.6 TB of memory (using --pad 1). Finally, Ewald sphere correction was applied to the final map, resulting in a resolution of 6.3 Å and a notable improvement in the map interpretability and FSC curve (**Supplementary Figure 4**).

### Structural analysis of Jyvaskylavirus

Using AlphaFold3 we generated an initial atomic model of the MCP corresponding to the ORF184 identified in this study by sequence comparison with other *Marseillaviridae* (**Abramson et al 2024**). The best-ranked predicted model had a confidence level of 95 % based on the predicted local-distance difference test (pLDDT), which ranged from 28.7 to 98.9 % across the residues. The MCP model was manually fitted into the density corresponding to the pseudo-hexameric capsomer using COOT graphic software (**Emsley & Cowtan 2004**). The three molecules were rigid body refined into the 6.3 Å resolution map using PHENIX real-space refinement (**Afonine et al 2018**) leading to a CC_mask_ of 52 %.

To identify the penton protein located at the vertices as well as some of the additional visible densities corresponding to the scaffolding and cementing proteins, we used the following strategy. Owing that Jyvaskylavirus belongs to the clade B of the *Marseilleviridae* (current study) and that block-based derived cryo-EM maps at the 2-, 3- and 5-folds of Melbournevirus (clade A) at about 3.5 Å resolution have been recently made available in the Electron Microscopy Database (EMD-37188, -37189, -37190) we used the latest version of ModelAngelo software (v1.0.12) without any fasta sequence input to build and identify the likely models for the penton and ancillary proteins in the Melbournevirus densities **(Burton-Smith and Murata 2023**; **Jamali et al 2024**). The sequences of the resulting identified amino acids for the built fragments were then parsed using the hidden Markov model (HMM) routine in ModelAngelo against the list of ORFs of Jyvaskylavirus produced in this study (**Supplementary Figure 5**). This was performed with the expectation of high sequence homology between the two viruses, as indicated by our comparative sequence analysis (**Figure 2**). For the penton protein, ORF142 was identified, and similarly ORF36, 97, 153 and 121 for the ancillary under the five-fold and cap protein, respectively (**Supplementary Figure 10**). Then, the corresponding full 3D models were predicted using AlphaFold3 and fitted into the Melbournevirus and Jyvaskylavirus cryoEM density using the fit-into-map routine in ChimeraX together with the peripentonal capsomers (**Meng et al 2023**). To assess the metric of this fitting (**Supplementary Figure 7**), the 3.5 Å five-fold Melbournevirus block 3D density (EMDB-37190) was boxed around the pentameric assembly model and refined as a whole using rigid-body refinement in PHENIX, yielding a CC_mask_ of 57.3%. The same pentameric model was subsequently fitted into the 6.3 Å Jyvaskylavirus 3D cryo-EM density (previously boxed around the model), resulting in a lower CC_mask_ of 33%, consistent with the limited resolution of the capsid map and below regions. In the case of Melbourne virus, the corresponding ORFs were determined by using the Jyvaskylavirus ORFs as templates in blastp (https://blast.ncbi.nlm.nih.gov/Blast.cgi?PAGE=Proteins). The predicted models mentioned above were deposited in BioStudies under S-BSST1654.

## Data Availability

The Jyvaskylavirus genome has been deposited in GenBank under the accession number PQ284187 as part of BankIt submission 2862875. Raw stacks movies are uploaded to EMPIAR (EMPIAR-12466), linked to the EMDB accession code 51613 corresponding to the reconstructed Jyvaskylavirus cryo-EM map at 6.3 Å resolution. Predicted Jyvaskylavirus PDB models using ModelAngelo and Alphafold have been deposited at BioStudies (https://www.ebi.ac.uk/biostudies/) under the accession number S-BSST1654.

## Author contributions

G.M.dF.A, L.R.S. and N.G.A.A. conceived the project. G.M.dF.A. performed bioprospection experiments and virus isolation. G.M.dF.A. and ML performed the EM and HIM imaging. J.A., B.L.A. and J.S.A. were responsible for sequencing and bioinformatics analyses. I.A. performed all cryo-EM experiments, including sample vitrification, image processing, cryo-EM density analysis and implementation of ORFs identification and models fitting under the guidance of N.G.A.A. D.Z. collected cryo-EM data with support from I.A. and N.G.A.A. G.M.dF. L.R.S. and N.G.A.A. wrote the manuscript with input from all authors.

## Acknowledgements

We would like to thank Mr Petri Papponen (University of Jyvaskyla, Finland) for help with the electron microscopy, Msc. Marjaana Hassani (University of Jyvaskyla, Finland) for donating the composting sample used for Jyvaskylavirus isolation and the staff at the EM Platform at the CIC bioGUNE for infrastructural support and preliminary cryo-grid screening. Professor Bernard La Scola (Aix Marseille University, France) is thanked for donating the amoeba strains used and for his technical and intellectual input in this project. Yun Song at the UK’s national Electron Bio-imaging Centre (eBIC) at Diamond Light Source (DLS) is acknowledged for her assistance in setting up the SerialEM script. We are also grateful to DLS for access and support of the cryo-EM facilities at eBIC [under proposal CM30316], funded by the Wellcome Trust, MRC and BBRSC and to the Spanish Supercomputing Network (RES, Red Española de Supercomputación) for Picasso resources provided by the Supercomputing and Bioinnovation Center (Malaga University) to BCV-2022-3-0015.

This study was funded by the Research Council of Finland (grant #346772), the Centre for New Antibacterial Strategies (CANS) at the Arctic University of Norway (project ID #2520855), the Spanish Ministry of Science, Innovation and Universities, and the State Research Agency (AEI) for project PID2021-126130OB-I00 (NGAA) and the Severo Ochoa Excellence Accreditation to CIC bioGUNE (CEX2021-001136-S), as well as by the Conselho Nacional de Desenvolvimento Científico e Tecnológico (CNPq), the Coordenação de Aperfeiçoamento de Pessoal de Nível Superior (CAPES), the Fundação de Amparo à Pesquisa do Estado de Minas Gerais (FAPEMIG), and the Pró-Reitorias de Pesquisa e Pós-Graduação da UFMG (PRPG-UFMG). JSA is a CNPq researcher. The publication charges for this article have been funded by a grant from the publication fund of UiT The Arctic University of Norway.

## SUPPLEMENTARY MATERIAL LIST

**Supplementary text (docx)**: Preliminary data about isolation of Finnish giant viruses using a locally isolated amoeba host; Jyvaskylavirus growth, chloroform sensitivity and stability.

**Supplementary Figure 1:** Details of our preliminary screening for Finnish giant viruses’ effort.

**Supplementary Figure 2:** Jyvaskylavirus growth curve, sensitivity to chloroform and stability.

**Supplementary Figure 3:** Phylogenetic tree reconstructed using Major Capsid Protein (MCP) amino acid sequences belonging to viruses from *Marseilleviridae* family.

**Supplementary Figure 4:** Fourier Shell Correlation curve of the whole Jyvaskylavirus.

**Supplementary Figure 5:** Schematic workflow for ORF identification and model prediction

**Supplementary Figure 6:** Fitting of individual into the corresponding densities of the 5-fold block Melbourne virus map

**Supplementary Figure 7:** Fitting of all Jyvaskylavirus protein models into the 5-fold block Melbourne virus map

**Supplementary Figure 8:** Fitting of Jyvaskylavirus ORF119 protein model into the 2-fold block Melbourne virus map

**Supplementary Figure 9:** A side-by-side comparison of the predicted Jyvaskylavirus ORF142 with experimentally derived penton proteins of the lipid-containing bacteriophage PRD1 and the archaeal virus Haloarcula California icosahedral virus 1, represented as cartoon tube models color-coded from blue to red in a rainbow gradient from the N-terminal to the C-terminal, with the corresponding PDB ID codes in parentheses.

**Supplementary Figure 10:** Sequence alignment of selected Jyvaskylavirus minor capsid protein sequences (ORF036, ORF097, and ORF153) with residues modeled into the Melbournevirus density map at ∼3.5 Å resolution using ModelAngelo, visualized with the ESPRIPT software (https://espript.ibcp.fr/ESPript/ESPript/).

**Supplementary Table 1 (xls):** Details of the samples used and isolation results.

**Supplementary Table 2 (xls):** Jyvaskylavirus annotation.

**Supplementary Table 3 (docx):** Grid vitrification parameters and cryo-EM data collection.

**Supplementary Table 4 (docx):** Identified Melbournevirus ORFs through Jyvaskylavirus ORFs.

