## Supplementary material for "Genomic and structural insights into Jyvaskylavirus, the first giant virus isolated from Finland"

**Supplementary text**

**Isolation of Finnish giant viruses using a locally isolated amoeba host.**

During the summer 2019 we isolated a native amoeba from Finland. Amoeba isolation attempts were made by adding a drop of water from different sources to a plate of non-nutrient (NN) agar covered with 100 µl of dead *Escherichia coli* cells. The plates were checked daily for the presence of amoebal-like growth. When cells were seen, a block of agar was cut and added upside down to a fresh NN-agar plate containing dead *E. coli*. Plates were then checked daily for the visualization of amoebal cells spreading on the agar from below the agar block. Images of the amoeba isolation process can be seen in **Supplementary Figure 1A-C.** The non-nutrient agar recipe used was 120 mg NaCl, 4 mg MgSO_4_ · 7H_2_O, 4 mg CaCl_2_ · 2H_2_O, 142 mg Na_2_HPO_4_, 136 mg KH_2_PO_4_ and 15g agar in one litre of water (**Thomas et al 2006**). Dead *E. coli* cells were prepared by autoclavation of a turbid overnight culture.

For isolating the amoeba ten freshwater samples were tested. These samples consisted of Jyvasjarvi lake water, water from fish farms, aquarium water and outlet water from the University of Jyväskylä aquarium facility. All samples had amoeba-like cells growing inside the initial drop. Four samples had amoebal-like cells spreading on the plate from the agar block cut. Of these, one was fast-growing while the other three were growing slow in the following passages. We chose the fast-growing isolate, originated from the University of Jyväskylä aquarium facility, and used it to screen for giant viruses in samples collected from Jyväskylä. We later discovered that this amoebal isolate carried a symbiotic bacterium, making it impossible to grow the amoeba in liquid media. Any attempt resulted in bacterial contamination of the media. Passing the amoeba in antibiotic containing media ended up killing the bacteria and the amoeba at the same time. Encystment of this species was not observed, and preservation by freezing was not effective, meaning that it became impossible to make stocks for future use.

Therefore, further isolation attempts for giant viruses using this amoeba isolate were made in non-nutrient agar plates to avoid the problem with liquid media contamination. Fresh amoeba cultures collected from a plate were mixed with the samples to be tested (1:1) and added as drops to a new non-nutrient agar plate covered with dead *E.coli* cells. Appearance of cytopathic effect (CPE) led to the harvesting of the sick cells and evaluation of viral-like particles through electron microscopy. Examples of a control culture and two cultures with CPE can be seen in **Supplementary Figure 1D-F**. CPE in this system resulted in alterations of cell morphology and of patterns of cell distribution on the plate, differing from the typical cell lysis seen in liquid cultures.

Electron microscopy revealed distinct virus-like particles were in these enrichment samples (**Supplementary Figure 1G-K**). However, we were unable to grow these using co-cultures with the original host on agar plates and these samples did not result in CPE when we inoculated them in the other amoebal species used in this study (*A. castellanii*, *A. polyphaga* and *V. vermiformis*). Considering that we were unable to grow these putative viruses and consequently could not further characterize them, we show this data here as preliminary evidence of a high diversity of unknown giant viruses in Finland. Host-range and the use of additional locally isolated amoebas should be considered in further studies.

**Jyvaskylavirus growth, chloroform sensitivity and stability**

Jyvaskylavirus grows fast in *A. castellanii* cells (**Supplementary Figure 2A**). It causes CPE in *V. vermiformis* in high concentrations only (10^10^ TCID50/ml), and in high to middle concentrations in *A. polyphaga* (10^10^ to 10^6^ TCID50/ml). Electron microscopy revealed a peculiar double-capsid structure with an indication of a lipid layer in between, so we decided to test whether chloroform would affect the virion stability. Exposure to 10% chloroform for ten minutes reduced the viral titer in around six logarithmic orders, indicating presence of a lipid moiety in the capsid (**Supplementary Figure 2B**). A stability test was made by leaving viral aliquots at different temperatures for over 100 days. Viruses were recovered from all temperatures including 37 °C, showing that the virus is stable for long periods without its host presence (**Supplementary Figure 2C**).

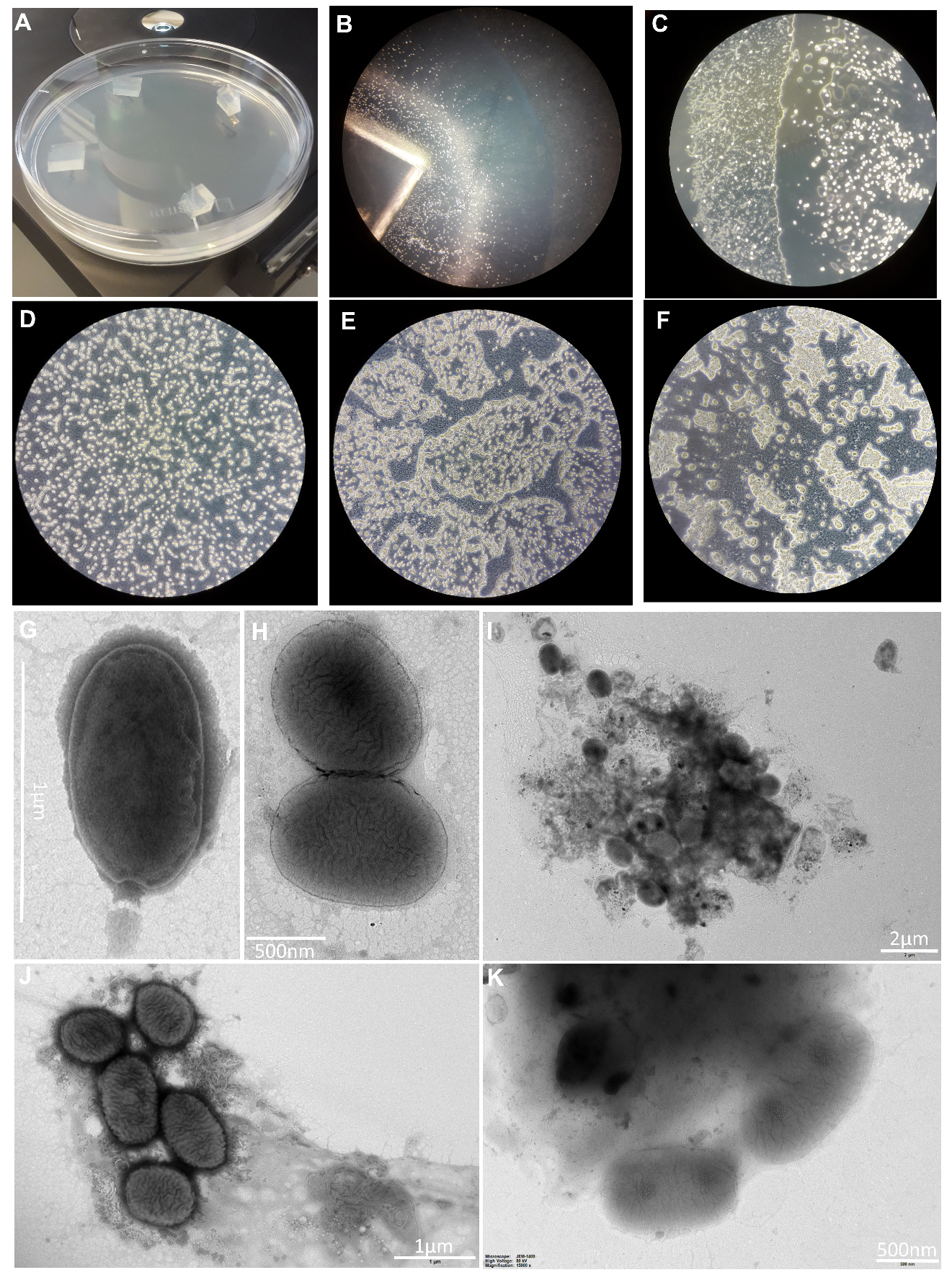

**Supplementary Figure 1**: Details of our preliminary screening for Finnish giant viruses effort. A) Non-nutrient agar plate used for amoebal isolation. B) Amoebal-like cells spreading from a block of agar. C) Amoebal-like cells growing on top of a non-nutrient agar plate. D) Control culture from an isolation attempt. E-F) Cytopathic effect seen in two different samples. G-K) Viral-like particles seen by electron microscopy.

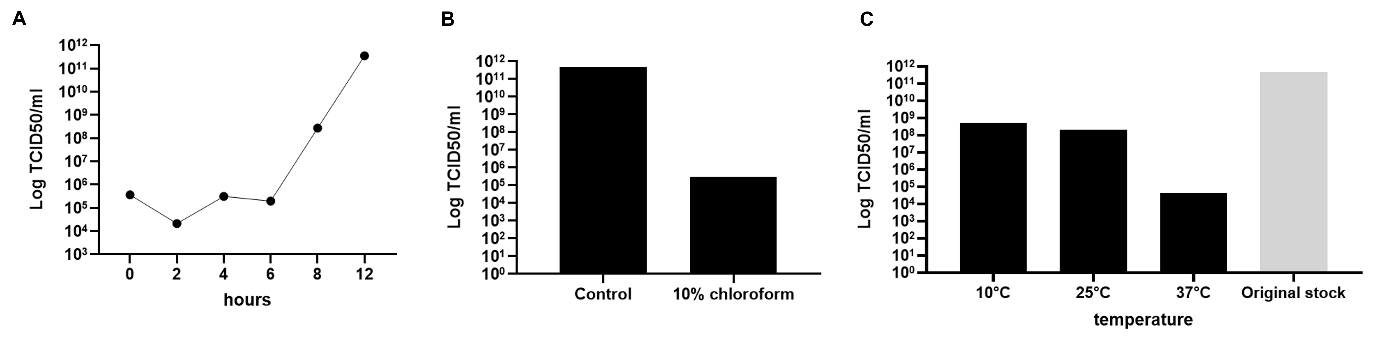

**Supplementary Figure 2:** Jyvaskylavirus growth curve, sensitivity to chloroform and stability. A) Growth curve in *A. castellanii* cells. B) Jyvaskylavirus sensitivity to chloroform exposure. C) Jyvaskylavirus stability after storage for 109 days in different temperatures. The original titer at day zero was 10^11^ TCID50/ml. Data shown in A and B were titrated in quadruplicates and in C in octuplicates.

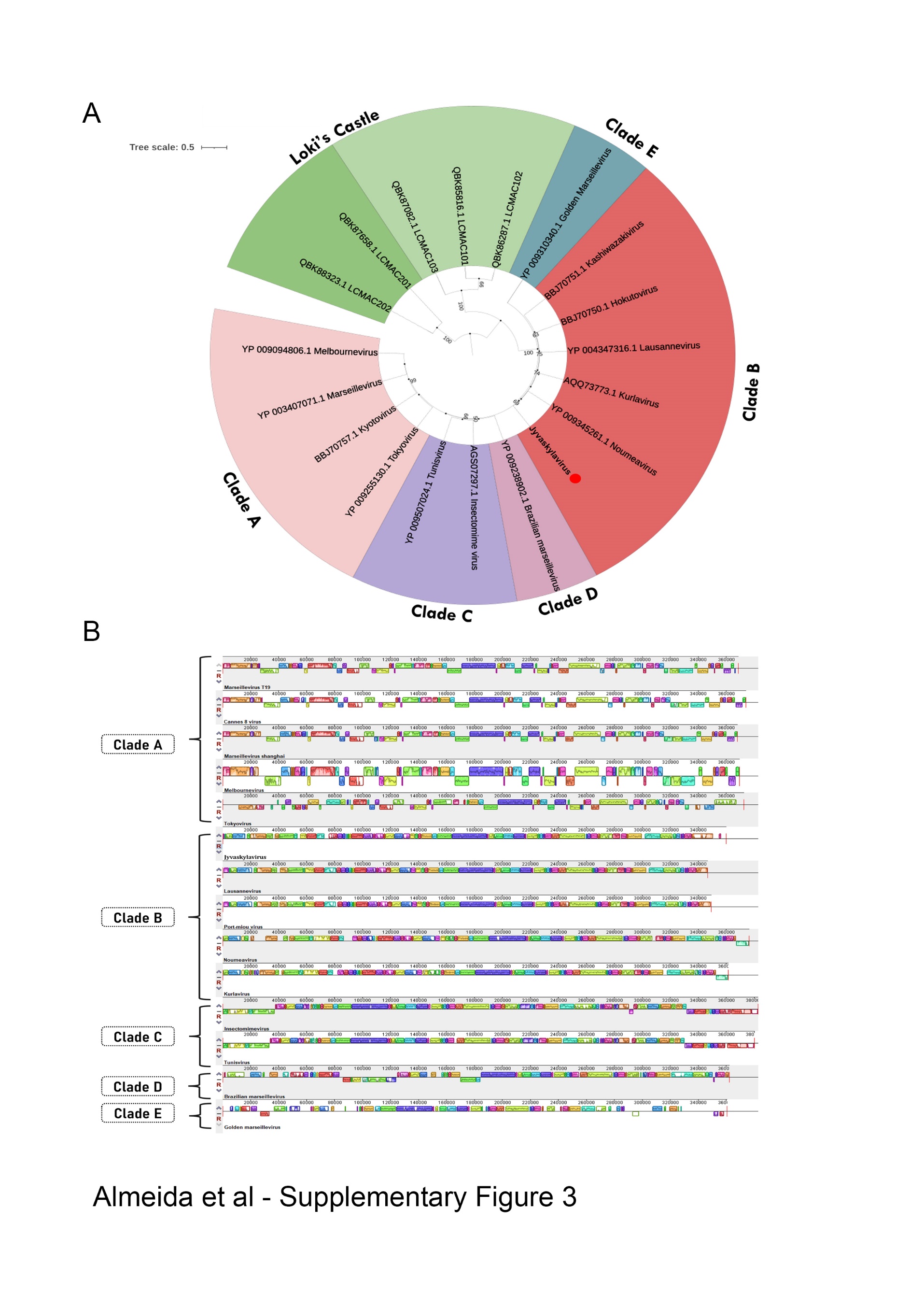

**Supplementary Figure 3:** A) Phylogenetic tree reconstructed using major capsid protein (MCP) amino acid sequences belonging to viruses from *Marseilleviridae* family. The Jyvaskylavirus sequence is highlighted in bold and indicated by a red circle. The alignment was performed with MUSCLE and the maximum-likelihood tree was reconstructed using IQtree software using ultrafast bootstrap (1000 replicates). The best-fit model selected using ModelFinder (implemented in IQtree) was rtREV+F+G4. Scale bar indicates the number of substitutions per site. B) Genome synteny scheme showing the MAUVE alignment of complete genome sequences belonging to marseilleviruses from five different lineages (A to E). Similar genome blocks are coded by the same color in each sequence. Blocks represented above de x-axis are in the forward strand whereas blocks represented below the x-axis are in the reverse strand.

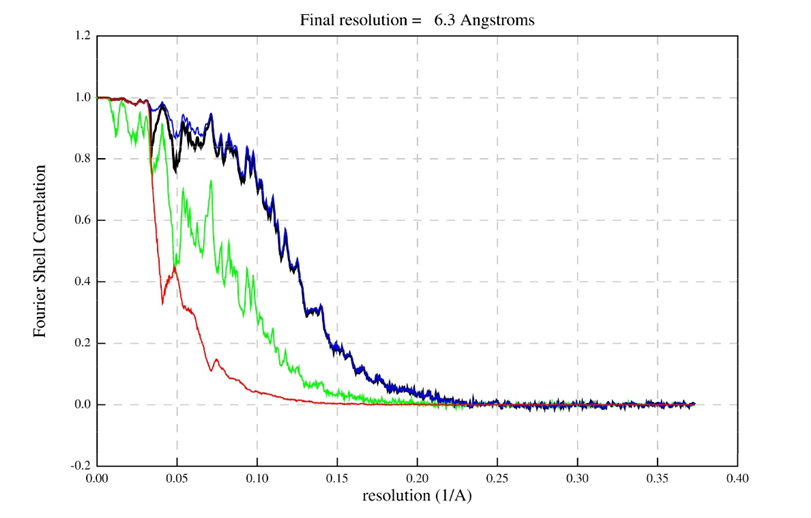

**Supplementary Figure 4:** A) Fourier Shell Correlation curve of the whole Jyvaskylavirus reporting a resolution of 6.3 Å at the 0.143 criterion (black line: corrected map); other colored curves indicate the following: blue line, masked maps; red line, phase-randomized masked maps; and green line, unmasked maps.

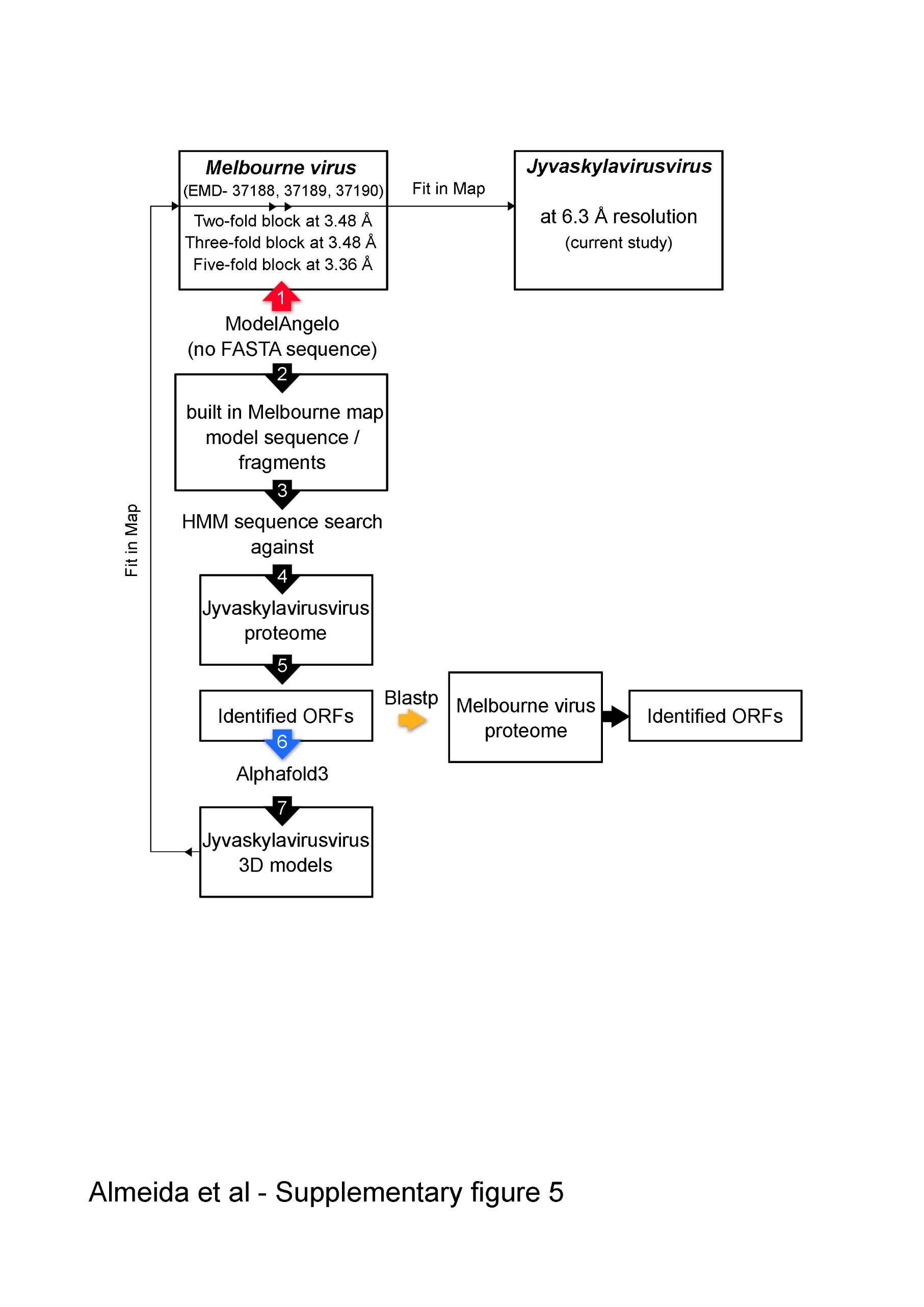

**Supplementary Figure 5:** Schematic workflow illustrating the identification of additional ORFs corresponding to ancillary proteins using an integrative approach combining ModelAngelo, Alphafold3 and ChimeraX software. Large, numbered arrows indicate the sequences of key steps in the workflow: the red arrow marks the use ModelAngelo to build polypeptide chains in the Melbournevirus maps; the blue arrow indicates the step of predicting the fold of the identified Jyvaskylavirus ORFs; and the orange arrow marks the step in which the sequences of the Jyvaskylavirus proteins were queried against the reference Melbournevirus isolate 1 (NCBI Reference Sequence: NC_025412.1).

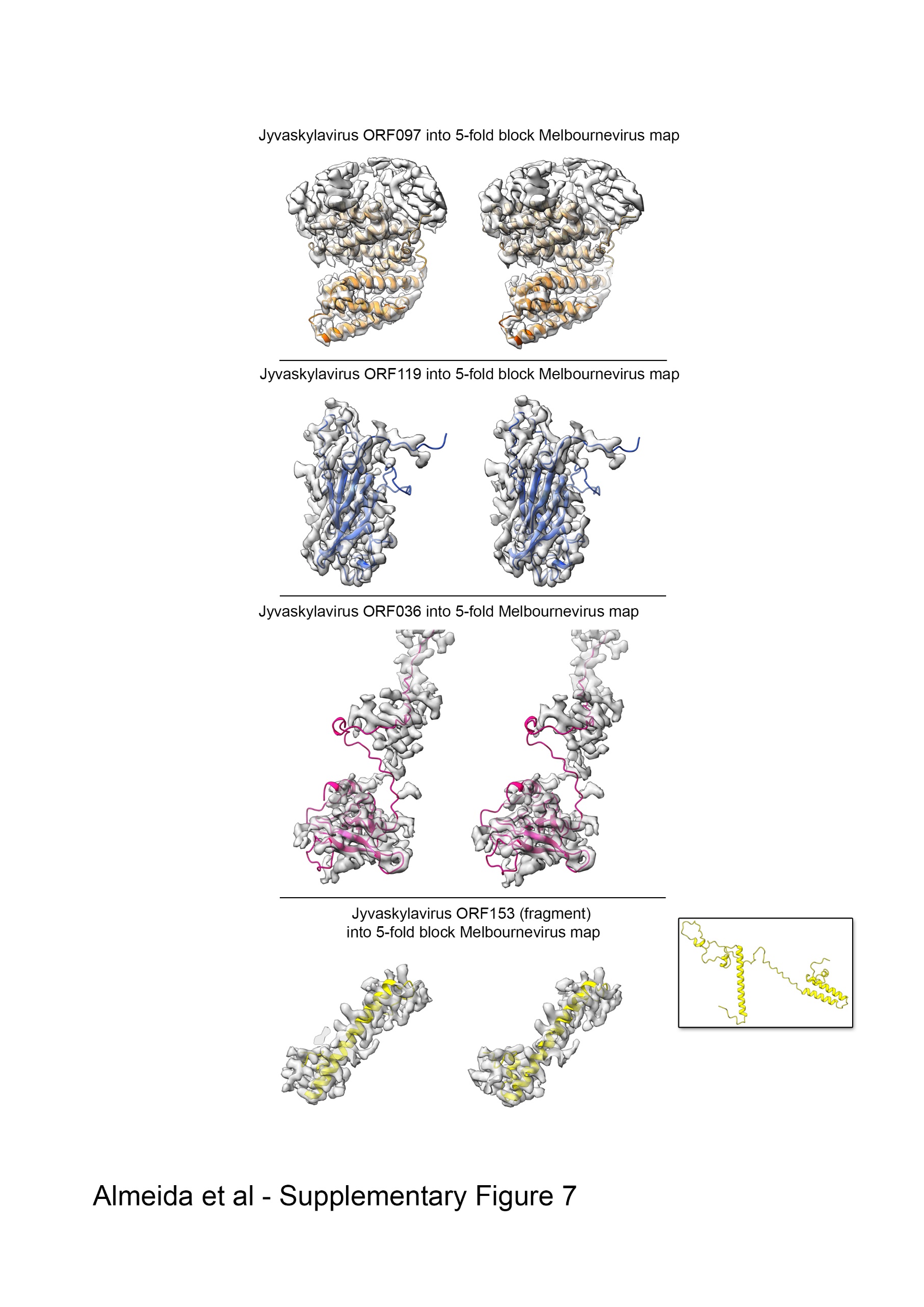
**Supplementary Figure 6:** Stereo-view (cross-eye) of the identified Jyvaskylavirus ORFs corresponding to the ancillary proteins beneath the capsid shell fitted into the cryo-EM Melbournevirus five-fold block reconstructed density (EMD-37190) shown in semi-transparent white; a B-factor of -5 Å^2^ and resolution cutoff to 4 Å have been applied to the original map for clarity. ORF097 forms a clear dimer while only a helical fragment of ORF153 has been depicted (the inset shows the full prediction). While the predicted core folds of the individual AlphaFold3 models align reasonably well with the density map, the terminal ends, predicted to be flexible, do not fit within the density.

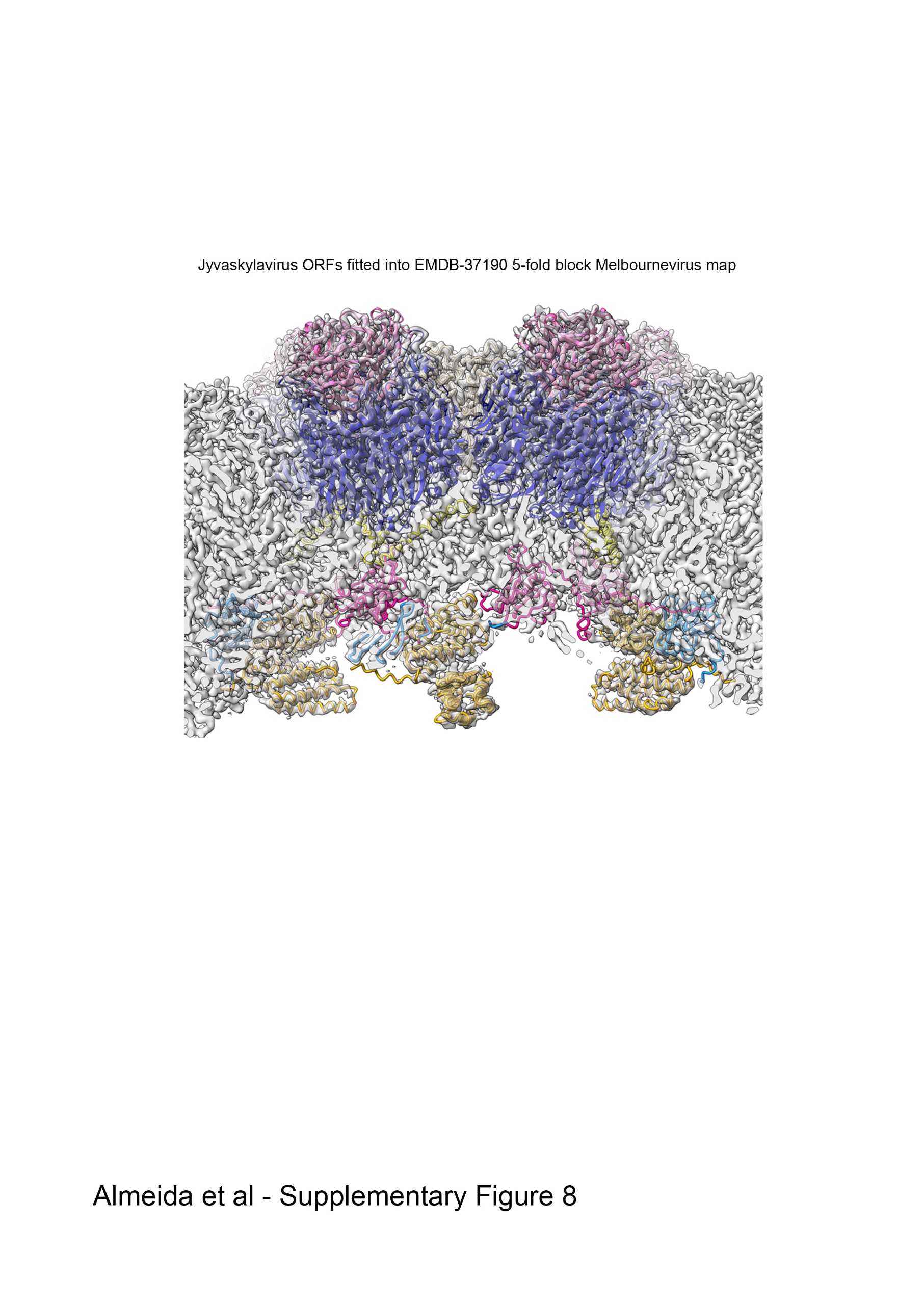

**Supplementary Figure 7:** Melbournevirus five-fold block reconstructed density (EMD-37190) in semi-transparent white (a B-factor of -5 Å^2^ and resolution cutoff to 4 Å have been applied to the original map for clarity), with the predicted Jyvaskylavirus 3D models of the major capsid protein (ORF184) fitted into the density, composing the peripentonal pseudo-hexameric capsomers represented as spheres in medium blue and capped at the top by ORF121 trimers (shown as pink spheres); five-copies of the penton protein (ORF142), represented as carton tubes in light-brown color plug the vertex. Below, different pentasymmetron protein components are represented as cartoon tubes, corresponding to a fragment of ORF153 (yellow), ORF97 (as dimer, color-coded in goldenrod and orange), ORF119 (dodger blue), and ORF36 (hot pink); not all the density below the vertices is accounted for with the fitted models. Boxing the map around the pentameric assembly atomic model and performing rigid-body fitting yielded a CC_mask_ of 57.3%

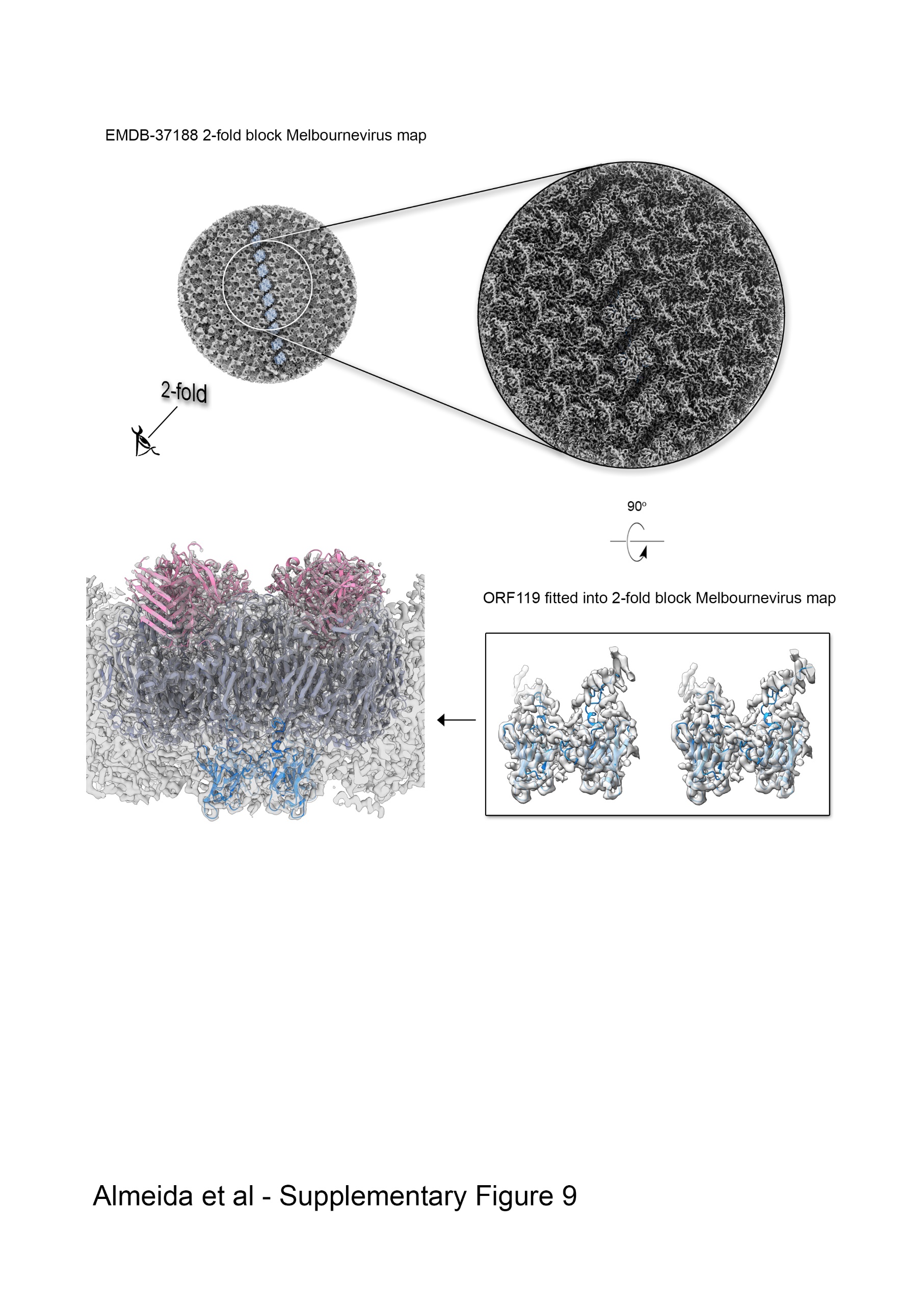

**Supplementary Figure 8:** Top left, view of the 2-fold block cryo-EM density reconstruction of Melbourne virus (Gaussian filtered for clarity and rendered as a light grey surface) as seen from within the virion along the 2-fold icosahedral symmetry axis; densities along the edge of two adjacent trisymmetron are colored in dodger-blue, similar to Figure 7A (top left). Top right, enlarged inset corresponding to the region marked by a white circle on the left; this shows the original density with an applied B-factor of -5 Å² and a resolution cutoff of 4 Å for clarity, revealing details of the secondary structure elements of different ancillary proteins. Bottom right, stereoview (cross-eye) of the ORF119 fitted into density showing the matching with the map with the exception of the terminal ends. Bottom left, fitting of two copies of capsomers displayed as in Supplementary figure 7 and viewed perpendicularly to the direction of the 2-fold axis showing the spatial organization between the copies of ORF119 and the capsomers.

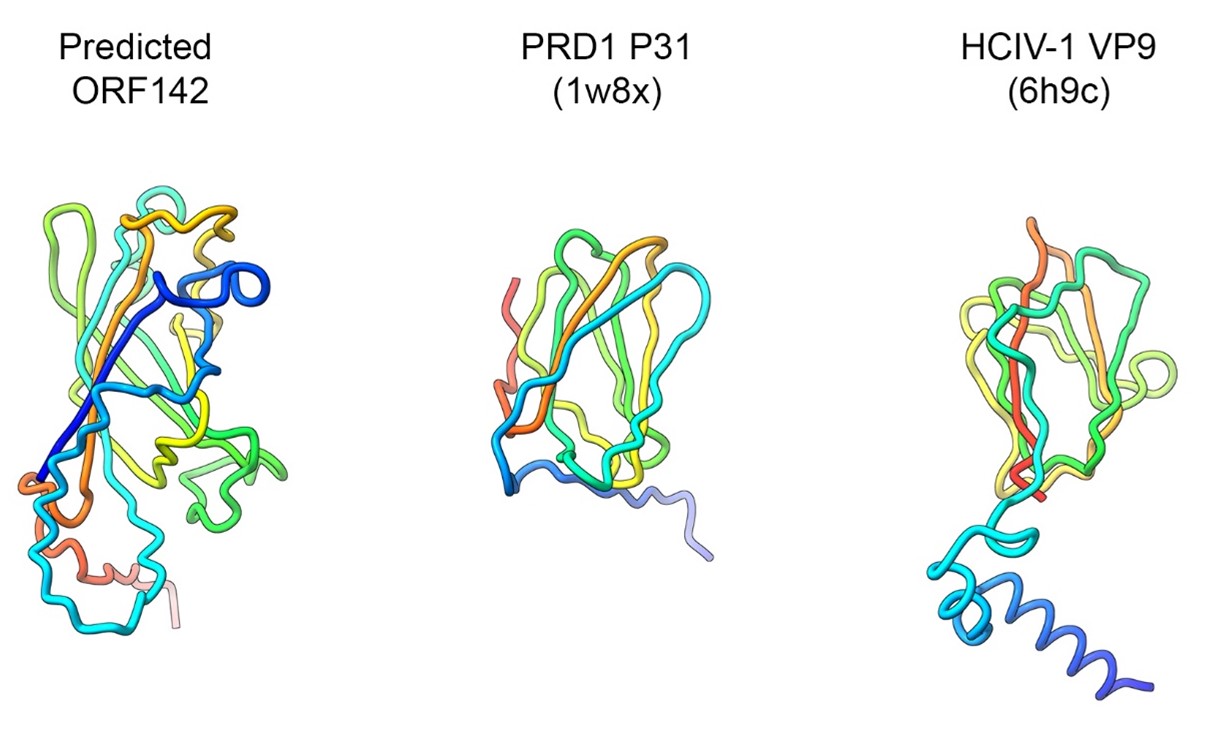

**Supplementary Figure 9:** A side-by-side comparison of the predicted Jyvaskylavirus ORF142 with experimentally derived penton proteins of the lipid-containing bacteriophage PRD1 and the archaeal virus *Haloarcula californiae* icosahedral virus 1, represented as cartoon tube models color-coded from blue to red in a rainbow gradient from the N-terminal to the C-terminal, with the corresponding PDB ID codes in parentheses.

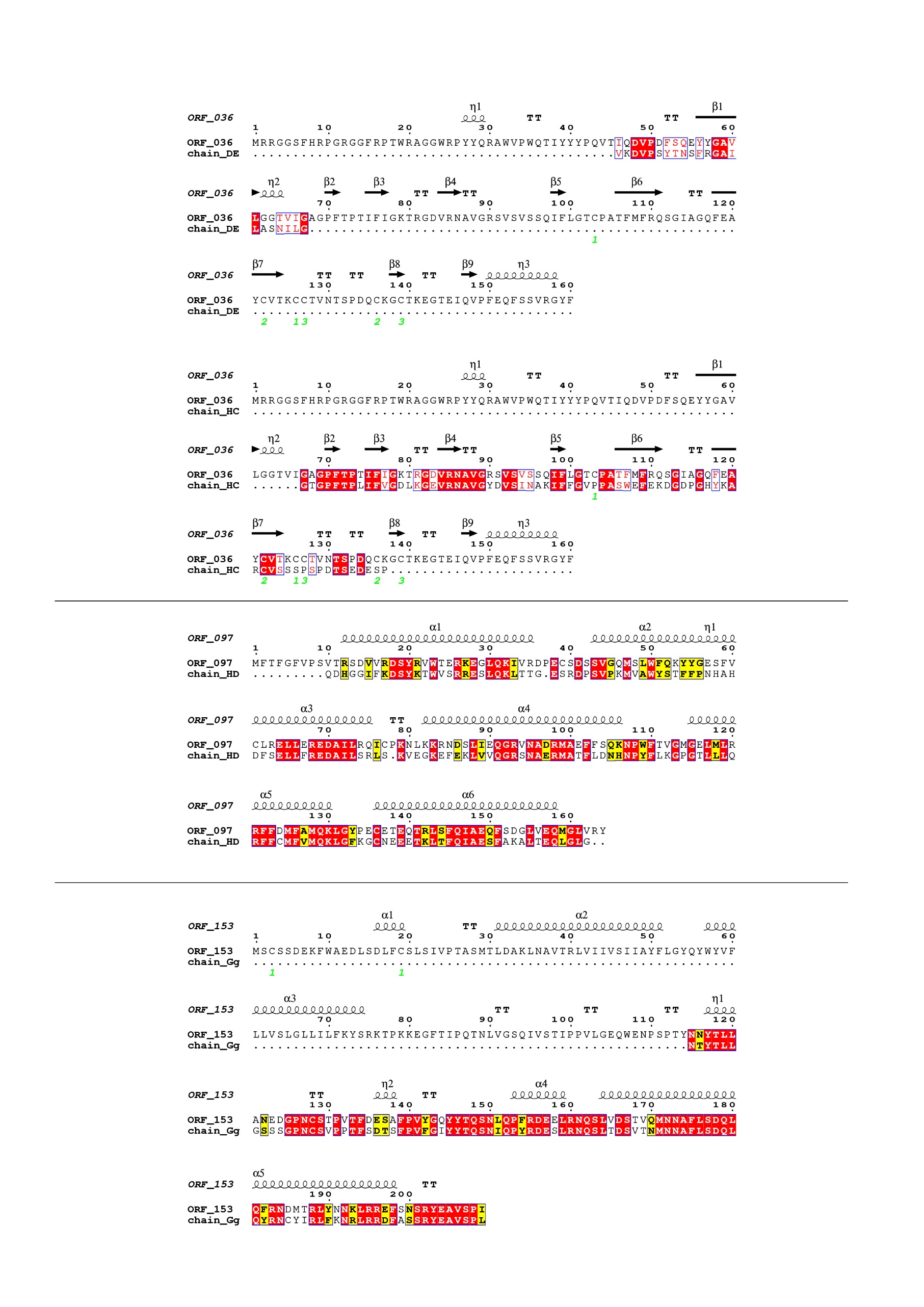

**Supplementary Figure 10:** Sequence alignment of selected Jyvaskylavirus minor capsid protein sequences (ORF036, ORF097, and ORF153) with residues modeled into the Melbournevirus density map at ~3.5 Å resolution using ModelAngelo, visualized with the ESPRIPT software (<https://espript.ibcp.fr/ESPript/ESPript/>). Strictly conserved residues are highlighted with red boxes and white characters, while similar residues are marked with yellow boxes and black characters. Secondary structure elements, as predicted by AlphaFold3, are shown above the alignment. Cysteine residues forming disulfide bonds in the predicted structure are indicated by green numbers below the alignment. The chains modeled by ModelAngelo are labeled according to the specific fragments that the software could build within the density regions.

**Supplementary Table 3. Grid vitrification parameters and cryo-EM data collection**

| **Data collection** | **1** | **2** | **3** | **4** |
| --- | --- | --- | --- | --- |
| *Session code @ eBIC* | bi23872-21 | cm30316-1 | cm30316-2 | cm30316-3 |
| *Date* | Jul 2021 | Dec 2021 | Jan 2022 | Mar 2022 |
| *ThermoFischer Krios 300 kV at eBIC* | Krios IV | Krios II | | |
| *Pixel size at specimen (Å/pix)* | 1.34 | 1.35 | | |
| *SuperRes Bin* | 1 | 2 | | |
| *Collection software* | EPU | Serial EM + ArbitrEM * | TOMO5  2 tilts: [0º , 5º] † | Serial EM |
| *Magnification* | ×64,000 | | | |
| *Detector* | BioQuantum K3 | | | |
| **Data collection & dose parameters** | | | | |
| *N. of Frames/Fractions* | 40 | 44 | 45 | 43 |
| *Electron Fluence*  *(e^-^/Å^2^)* | 40 | 44.5 | 45 | 45.77 |
| *Electron Flux (e^2^/pix/sec)* | 14.7 | 4.8 – 11.0  (due to ice thickness) | 16.545 | 8 |
| *Exposure time (sec)* | 5 | 5.5 | 5 | 10.4 |
| *Defocus range (µm)* | -1.2 to -2.3 | -1 to -2.6 (0.2 step) | -0.6 to -3 (0.3 step) | -1.2 to -2.2 (0.2 step) |
| **Sample preparation** | | | | |
| *Grids* | Quantifoil Cu R2/2 300 mesh | Quantifoil Cu R2/2 300 mesh + Extra carbon layer on top | Quantifoil Cu R2/2 300 mesh | Quantifoil Cu R2/1 300 mesh |
| *Vitrification equipment* | Vitrobot Mark IV (ThermoFischer) | Automatic Plunge Freezer EM2 (Leica) | Automatic Plunge Freezer EM2 (Leica) | Automatic Plunge Freezer EM2 (Leica) |
| *Vitrification settings* | - Sample droplet: 4 µL - Incubation time: 30 sec - Blot time: 2 sec - Offset value: -2, -3 - Humidity: > 95 % - Temperature: ≈ 4 ºC | - Sample droplet: 3 µL - Incubation time: 30 sec - Blot time: 0.9 sec - Humidity: > 90 % - Temperature: ≈ 8 ºC | - Sample droplet: 3 µL - Incubation time: 30 sec - Blot time: 1.5 sec - Humidity: > 90 % - Temperature: ≈ 8 ºC | - Sample droplet: 3 µL - Incubation time: 30 sec - Blot time: 1.5 sec - Humidity: > 90 % - Temperature: ≈ 8 ºC |
| **Preprocessing stats** | | | | |
| *N. of movies* | 14,644 | 288 | 383 | 1,955 |
| *Extracted particles* | 1,608 | 282 | 455 | 2,016 |
| *Useful particles* | 1,173 | 277 | 437 | 1,855 |
| *Particle contributing to the final map* | 3,742 | | | |
| ***** ArbitrEM software: <https://github.com/kyledent/ArbitrEM>  † The TOMO5 version used for that data collection did not allow to collect a single tilt fraction. Thus, [0º] and [-5º] tilts were collected per view, and then the [-5º] fractions were discarded. | | | | |

**Supplementary Table 4. Identified Melbournevirus ORFs through Jyvaskylavirus ORFs**

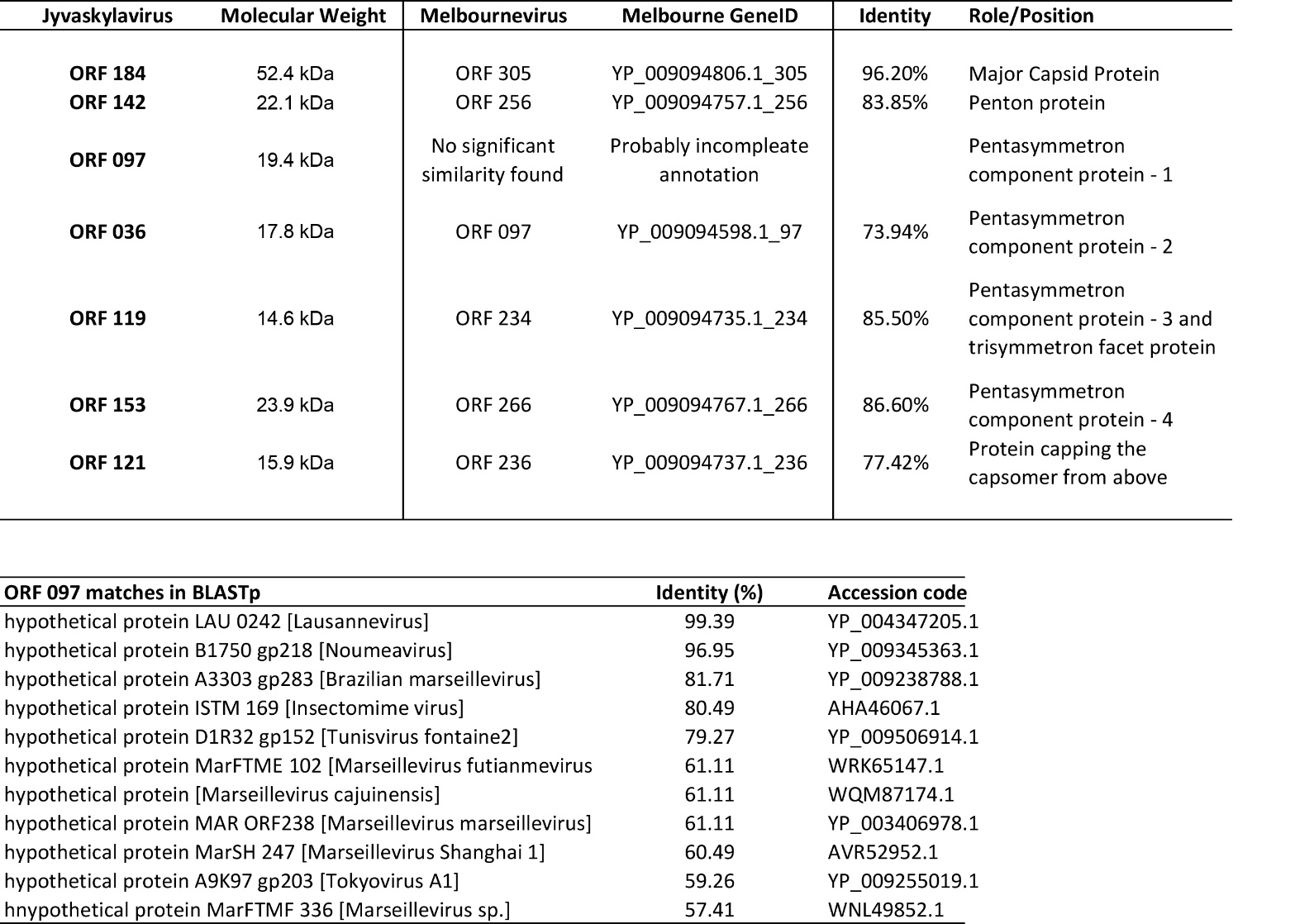

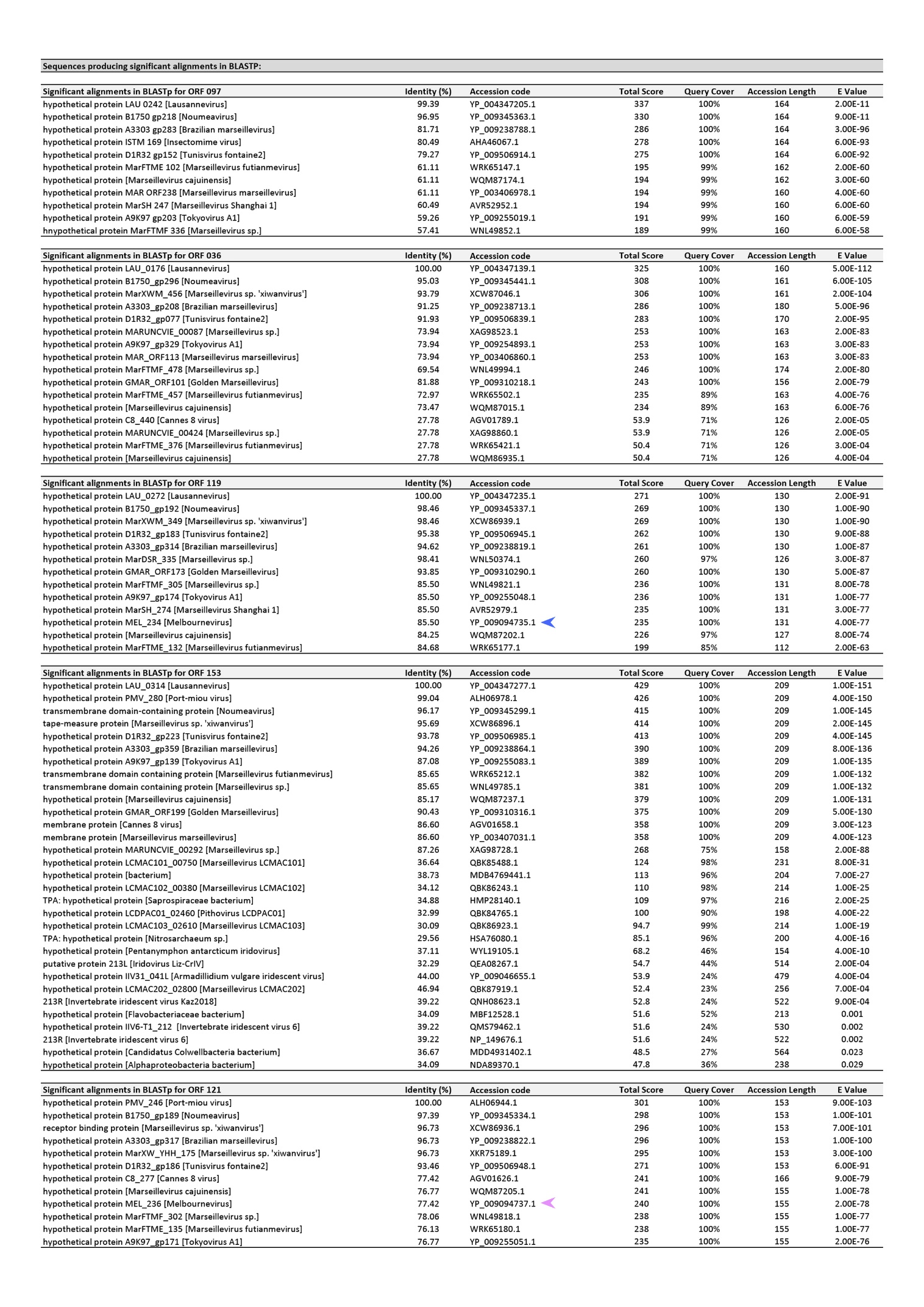
**Supplementary Table 5. Homologous sequences retrieved by BLASTp when querying** **Jyvaskylavirus minor capsid protein sequences**

**the colored arrows mark the Melbournevirus hits*

**Supplementary material references**

Thomas, V., Herrera-Rimann, K., Blanc, D. S., & Greub, G. (2006). Biodiversity of amoebae and amoeba-resisting bacteria in a hospital water network. Applied and Environmental Microbiology. https://doi.org/10.1128/AEM.72.4.2428-2438.2006
